# HYENA detects oncogenes activated by distal enhancers in cancer

**DOI:** 10.1101/2023.01.09.523321

**Authors:** Anqi Yu, Ali E. Yesilkanal, Ashish Thakur, Fan Wang, Yang Yang, William Phillips, Xiaoyang Wu, Alexander Muir, Xin He, Francois Spitz, Lixing Yang

**Affiliations:** Ben May Department for Cancer Research, University of Chicago, Chicago IL, USA; Department of Human Genetics, University of Chicago, Chicago IL, USA; University of Chicago Comprehensive Cancer Center, Chicago, IL, USA

## Abstract

Somatic structural variations (SVs) in cancer can shuffle DNA content in the genome, relocate regulatory elements, and alter genome organization. Enhancer hijacking occurs when SVs relocate distal enhancers to activate proto-oncogenes. However, most enhancer hijacking studies have only focused on protein-coding genes. Here, we develop a computational algorithm “HYENA” to identify candidate oncogenes (both protein-coding and non-coding) activated by enhancer hijacking based on tumor whole-genome and transcriptome sequencing data. HYENA detects genes whose elevated expression is associated with somatic SVs by using a rank-based regression model. We systematically analyze 1,146 tumors across 25 types of adult tumors and identify a total of 108 candidate oncogenes including many non-coding genes. A long non-coding RNA *TOB1-AS1* is activated by various types of SVs in 10% of pancreatic cancers through altered 3-dimensional genome structure. We find that high expression of *TOB1-AS1* can promote cell invasion and metastasis. Our study highlights the contribution of genetic alterations in non-coding regions to tumorigenesis and tumor progression.

## Introduction

At mega-base-pair scale, linear DNA is organized into topologically associating domains (TADs) ^1^, and gene expression is regulated by DNA and protein interactions governed by 3D genome organization. Enhancer-promoter interactions are mostly confined within TADs ^2–4^. Non-coding somatic single nucleotide variants (SNVs) in promoters and enhancers have been linked to transcriptional changes in nearby genes and tumorigenesis ^5^. Structural variations (SVs), including deletions, duplications, inversions, and translocations, can dramatically change TAD organization and gene regulation ^6^ and subsequently contribute to tumorigenesis. Previously, we discovered that *TERT* is frequently activated in chromophobe renal cell carcinoma by relocation of distal enhancers ^7^, a mechanism referred to as enhancer hijacking (**Fig. 1A**). In fact, many oncogenes, such as *BCL2* ^8^, *MYC* ^9^, *TAL1* ^10^, *MECOM/EVI1* ^11^, *GFI1* ^12^, *IGF2* ^13^, *PRDM6* ^14^, and *CHD4* ^15^, can be activated through this mechanism. These examples demonstrate that genomic architecture plays an important role in cancer pathogenesis. However, the vast majority of the known enhancer hijacking target oncogenes are protein-coding genes, and few non-coding genes have been reported to promote diseases through enhancer hijacking. Here, we refer to non-coding genes as all genes that are not protein-coding. They include long non-coding RNAs (lncRNAs), pseudogenes, and other small RNAs such as microRNAs, small nuclear RNAs (snRNAs), small nucleolar RNAs (snoRNAs), etc. They are known to play important roles in many biological processes ^16^, and some are known to drive tumorigenesis ^17,18^. In this study, we will focus on identifying oncogenes, including oncogenic non-coding genes, activated by enhancer hijacking.

**Figure 1.**
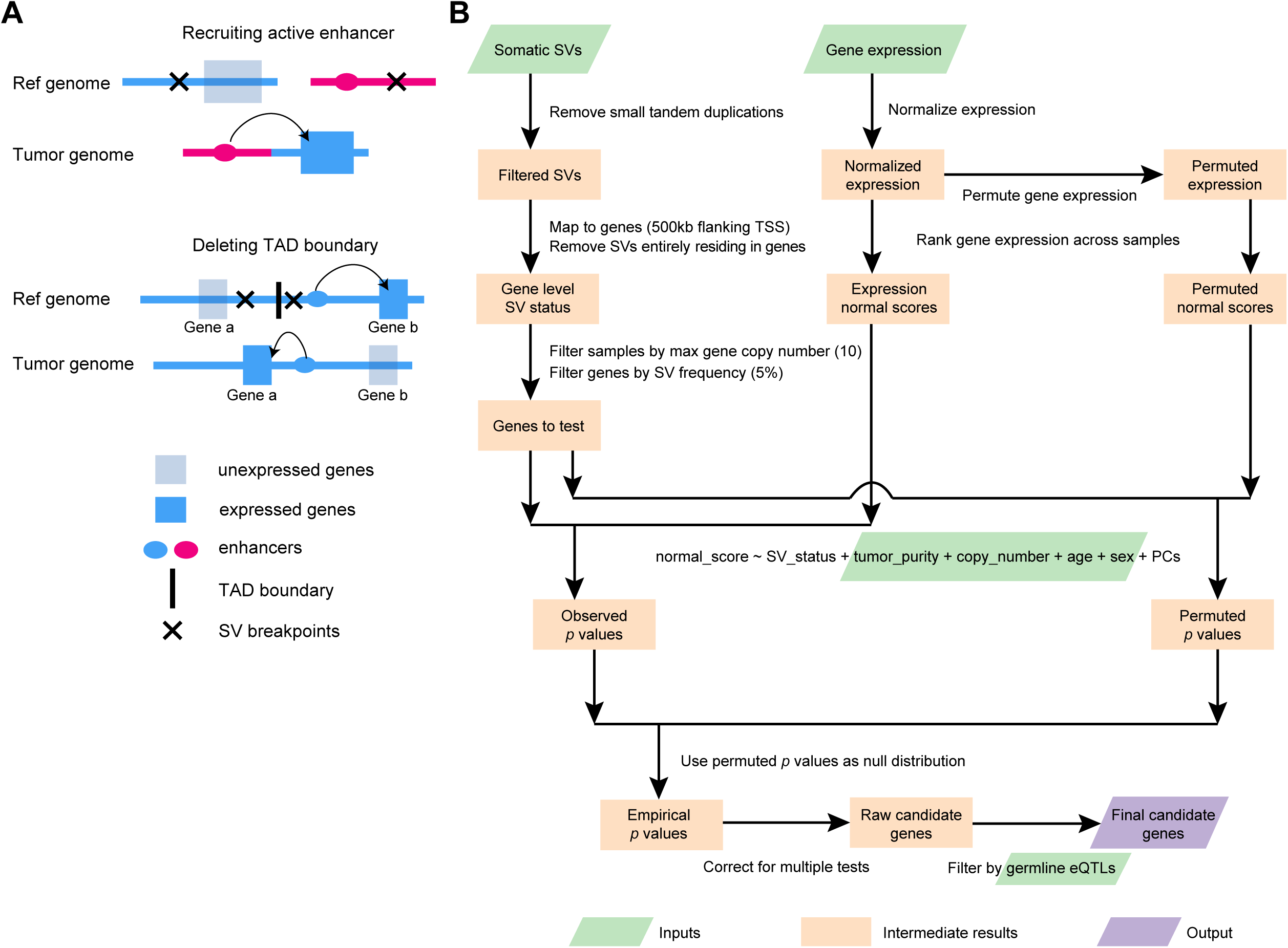
Outline of enhancer hijacking and HYENA algorithm. **A**, Mechanisms of gene activation by SVs. SVs can activate genes by recruiting distal active enhancers (top panel) and by removing TAD boundaries and forming de novo enhancer-promoter interactions (bottom panel). **B**, HYENA workflow. Green and purple boxes denote input and output files, respectively. Orange boxes denote intermediate steps. Numbers in parentheses represent default values of HYENA.

Several existing algorithms can detect enhancer hijacking target genes based on patient cohorts, such as CESAM ^13^ and PANGEA ^15^. These two algorithms implemented linear regression and elastic net model (also based on linear regression) to associate elevated gene expression with nearby SVs, respectively. PANGEA also considers the effects of somatic SNVs on gene expression. However, a major drawback of these algorithms is that linear regression is quite sensitive to outliers. Outliers are very common in gene expression data from cancer samples and can seriously impair the performances of these algorithms. In addition, CESAM is optimized for microarray data, while PANGEA depends on the annotation of tissue-specific promoter-enhancer pairs, which are not readily available for many tumor types. Cis-X ^19^ and NeoLoopFinder ^20^ can detect enhancer hijacking target genes based on individual samples. However, these tools have limitations in detectable genes and input data. Cis-X detects *cis*-activated genes based on allele-specific expression, which requires the genes to carry heterozygous SNVs. NeoLoopFinder takes Hi-C, Chromatin Interaction Analysis with Paired-End Tag (ChIA-PET), or similar data measuring chromatin interactions as input, which remain very limited. Furthermore, the identification of recurrent mutational events that result in oncogenic activation requires large patient cohorts. Therefore, tools that use whole-genome and transcriptome sequencing data, which are available at much larger sample sizes, would be more useful in identifying SV-driven oncogene activation. Finally, no non-coding oncogenes have been reported as enhancer hijacking targets by the above algorithms. A recent study on SVs altering gene expression in Pan-Cancer Analysis of Whole Genomes (PCAWG) samples ^21^ only considered protein-coding genes but not non-coding genes.

Here, we developed <u>Hi</u>jacking of <u>En</u>hancer <u>A</u>ctivity (HYENA) using normal-score regression and permutation test to detect candidate enhancer hijacking genes (both protein-coding and non-coding genes) based on tumor whole-genome and transcriptome sequencing data from patient cohorts. Among the 108 putative oncogenes detected by HYENA, we studied the oncogenic functions of a lncRNA, *TOB1-AS1*, and demonstrated that it is a regulator of cancer cell invasion in vitro and tumor metastasis in vivo.

## Methods

### Datasets

This study used data generated by the Pan-Cancer Analysis of Whole Genomes (PCAWG). We limited our study to a total of 1,146 tumor samples for which both whole-genome sequencing (WGS) and RNA-Seq data were available. The data set was composed of cancers from 25 tumor types including 23 bladder urothelial cancers (BLCA), 88 breast cancers (BRCA), 20 cervical squamous cell carcinomas (CESC), 68 chronic lymphocytic leukemias (CLLE), 51 colorectal cancers (COAD/READ), 20 glioblastoma multiforme (GBM), 42 head and neck squamous cell carcinomas (HNSC), 43 chromophobe renal cell carcinomas (KICH), 37 renal clear cell carcinomas from the United States (KIRC), 31 renal papillary cell carcinomas (KIRP), 18 low-grade gliomas (LGG), 51 liver cancers from United States (LIHC), 67 liver cancers from Japan (LIRI), 37 lung adenocarcinomas (LUAD), 47 lung squamous cell carcinomas (LUSC), 95 malignant lymphomas (MALY), 80 ovarian cancers (OV), 74 pancreatic cancers (PACA), 19 prostate adenocarcinomas (PRAD), 49 renal clear cell carcinomas from European Union/France (RECA), 34 sarcomas (SARC), 34 skin cutaneous melanomas (SKCM), 29 stomach adenocarcinomas (STAD), 47 thyroid cancers (THCA), and 42 uterine corpus endometrial carcinomas (UCEC). More detailed information on the sample distribution and annotation can be found in **Supplementary Table S1**.

WGS and RNA-Seq data analysis of tumor and normal samples were performed by the PCAWG consortium as previously described ^21^. Somatic and germline SNVs, somatic copy number variations (CNVs), SVs, and tumor purity were detected by multiple algorithms and consensus calls were made. Genome coordinates were based on the hg19 reference genome and GENCODE v19 was used for gene annotation. Gene expression was quantified by HT-Seq (version 0.6.1p1) as fragments per kilobase of million mapped (FPKM). Clinical data such as donor age and sex were downloaded from the PCAWG data portal (https://dcc.icgc.org/pcawg). *TOB1* and *TOB1-AS1* expression data in CCLE pancreatic cancer cell lines were downloaded from DepMap Public 22Q2 version (https://depmap.org/portal/download/all/). Gene expression data of the Cancer Genome Atlas (TCGA) PAAD cohort (TCGA.PAAD.sampleMap/HiSeqV2_PANCAN) and International Cancer Genome Consortium (ICGC) PACA-CA cohort for 45 samples of which “analysis-id” were labeled as “RNA” were downloaded from Xena Data Hubs (https://xenabrowser.net/datapages/) and ICGC data portal (https://dcc.icgc.org/projects/PACA-CA) respectively.

Significant expression quantitative trait loci (eQTL)-gene pairs (v8) were downloaded from the Genotype-Tissue Expression (GTEx) data portal (https://gtexportal.org/home/datasets). Only those eQTLs that had a hg19 liftover variant ID were included in the analysis and hg38 variants without corresponding hg19 annotation were discarded.

The raw sequencing data for Hi-C and ATAC-Seq were available through NCBI Sequence Read Archive (SRA) with accession number PRJNA1036282. The raw sequencing data for mouse xenograft tumor RNA-Seq were available through NCBI SRA with accession number PRJNA1011356.

### HYENA algorithm

First, small tandem duplications (<10 kb) were discarded since they are unlikely to produce new promoter-enhancer interactions. The remaining SVs were mapped to the flanking regions (500 kb upstream and downstream of transcription start sites [TSSs]) of annotated genes. SVs that fall entirely within a gene body were also discarded. The SV status of each gene was defined by the presence or absence of SV breakpoints within the gene or its flanking regions for each tumor. The binary variable SV status was used in the normal-score regression model below. Only genes carrying SVs in at least 5% of samples carrying SVs were tested. For each gene, samples with that gene highly amplified (>10 copies) were removed from the regression model.

#### Gene expression normal scores

Gene expression quantifications (fragments per kilobase per million [FPKM]) were quantile normalized (FPKM-QN) using the *quantile.normalize()* function from the *preprocessCore* R package to enhance cross-sample comparison. For each gene, samples were ranked based on their expression values, the ranks were mapped to a standard normal distribution and the corresponding z scores were gene expression normal scores. Normal-score conversion forced the expression data into a Gaussian distribution, allowing for parametric comparisons between samples.

#### Normal-score regression

A generalized linear model was used to test associations between gene expression normal scores and SV status and control for confounding variables such as gene copy number, tumor sample purity, donor age, and sex. To capture unobserved variations in gene expression, the first n principal components (PCs) of the expression data were also included in the regression model, where n was determined as 10% of the sample size of the cohort and up to 20 if the sample size was more than 200. The regression model was as shown below:

Expression_normal_score ∼ sv_status + copy_number + purity + age + sex + PC_1_ + PC_2_ …+ PC_n_

For each gene, all PCs were tested for associations with the SV status of that gene, and those PCs that significantly correlate (Mann-Whitney test, *P*<0.05) with SV status were not used in regression. A similar strategy was used to detect eQTLs in normal tissues ^22^.

#### Calculating empirical P values and model selection

Gene expression data were permuted 1000 times by randomly shuffling expression values within the cohort. For tumor types with more than 10,000 genes to test (**Supplementary Table S1**), only 100 permutations were performed to reduce run time. The normal-score regression was performed in the same way on observed gene expression and permuted expression. *P* values for SV status from permuted expression were pooled as a null distribution. Then the *P* values for SV status from observed expression and the *P*-value null distribution were used to calculate empirical *P* values. One-sided *P* values were used since we were only interested in elevated gene expression. False discovery rates (FDRs) were calculated using the Benjamini-Hochberg procedure. Genes with FDR less than 0.1 were considered candidate genes. For example, in MALY, there were 1,863 genes reaching 5% SV frequency and 1,863 *P* values were obtained in each permutation. After 1000 permutations, 1,863,000 *P* values were generated and should represent the null distribution very well. Empirical *P* values were calculated using these 1,863,000 permuted *P* values.

The above empirical *P* value calculation and candidate gene detection were performed iteratively with no PCs and up to n PCs in the regression model. When different numbers of PCs were included in the model, the numbers of candidate genes varied. The regression model with the lowest number of PCs reaching 80% of the maximum number of candidate genes in all regression models tested was selected as the final model to avoid over fitting. For example, the sample size for PCAWG UCEC was 42; therefore, we tested from 0 to 4 PCs. Among these, the model including 4 PCs gave the highest number (4) of candidate genes. Therefore, the model including 4 PCs with 4 candidate genes was selected as the final model (**Supplementary Table S2**).

In our normal-score regression, we essentially attempt to model variations in gene expression. Including confounding factors will improve performance. Tumor purity, gene copy number, patient age, and sex are factors known to affect gene expression. Therefore, they were included in the regression model. Unobserved variations may include tumor subtype, tumor stage, patient ethnicity, smoking status, alcohol consumption, and other unknown factors that may alter gene expression. Since HYENA was designed for wide applications, we did not require users to provide information on tumor subtype, tumor stage, patient ethnicity, smoking status, alcohol consumption, etc. Principle component analysis is a linear decomposition of gene expression variations. Therefore, including PCs in a regression model was suitable for removing systematic variations and could better model the effects of SV status. However, some enhancer hijacking target genes are master transcription factors, such as *MYC*, and have a profound impact on the gene expression of multiple pathways. Hence, it is possible that some PCs capture the activities of transcription factors. If these transcription factors were activated by somatic SVs, the PCs would be correlated with SV status. Including these PCs would diminish our ability to detect the effects of SV status. Therefore, we excluded these PCs from the regression model.

#### Testing eQTL-SV associations

Known germline eQTLs from the matching tissues were obtained from GTEx (**Supplementary Table S3**). The associations between germline genotypes of eQTLs and SV status of the candidate genes in the PCAWG cohort were tested using a Chi-squared test. Genes with significant correlations (*P*<0.05) between their SV status and at least one eQTL were removed. The remaining genes were our final candidate enhancer-hijacking target genes.

### Benchmarking

Known enhancer hijacking target genes in PCAWG tumor types were selected to test the sensitivity of HYENA, CESAM and PANGEA. The genes included *MYC* in malignant lymphoma, *BCL2* in malignant lymphoma, *CCNE1* in stomach/gastric adenocarcinoma, *TERT* in chromophobe renal carcinoma, *IGF2* in colorectal cancer, *IGF2* in stomach/gastric adenocarcinoma, *IGF2BP3* in thyroid cancer, and *IRS4* in lung squamous cell carcinoma. The same SVs, CNVs, and SNVs were used as input for all three algorithms. For CESAM and PANGEA, upper-quantile normalized fragments per kilobase per million (FPKM-UQ) were normalized by tumor purity and gene copy number, and then used as gene expression inputs. CESAM was run using default parameters, and FDR of 0.1 was used to select significant genes. PANGEA requires predicted enhancer-promoter (EP) interactions based on ChIP-Seq and RNA-Seq data. The EP interactions were downloaded from EnhancerAtlas 2.0 (http://www.enhanceratlas.org/) (**Supplementary Table S4**). EP interactions from multiple cell lines of the same type were merged. PANGEA was run with default parameters as well and significant genes were provided by PANGEA (multiple testing adjusted *P* value <0.05). To test HYENA, CESAM, and PANGEA for false positives, 20 random gene expression datasets for malignant lymphoma and breast cancer were generated by randomly shuffling sample IDs in gene expression data. HYENA, CESAM, and PANGEA were run with random expressions in the same way as above.

### Predicting 3D genome organization

A 1 Mb sequence was extracted from the reference genome centered at each somatic SV breakpoint and was used as input for Akita ^23^ to predict the 3D genome organization. Two 500 kb sequences were merged according to the SV orientation to construct the sequence of the rearranged genome fragments. Akita was used to predict the genome organization for the rearranged sequence. High-resolution Micro-C data obtained from human H1-ESCs and HFF cells ^24^ were used to facilitate TAD annotation together with predicted genome organization. H3K27Ac and CCCTC-binding factor (CTCF) ChIP-Seq data from the PANC-1 cell line were downloaded from the ENCODE data portal (https://www.encodeproject.org/). SV breakpoints were provided to Orca ^25^ to predict 3D genome structures through its web interface (https://orca.zhoulab.io/).

### In situ Hi-C and ATAC-Seq

Ten million cells of Panc 10.05, PANC-1, PATU-8988S, and PATU-8988T cell lines were collected to construct Hi-C libraries ^26^. The Hi-C libraries were sequenced on Illumina NovaSeq X Plus platform with 1% phix. About 2 billion reads were obtained from Panc 10.05, PATU-8988S, and PATU-8988T, and 1 billion reads were obtained from PANC-1. The paired-end reads were aligned to chromosomes 1-22, X, Y and M by bwa-mem. SVs were identified by EagleC ^27^ at 5 kb, 10 kb and 50 kb resolutions. The non-redundant SVs in **Supplementary Table S5** were combined for the three resolutions. Chromatin loops were identified by NeoLoopFinder ^20^. A probability threshold of 0.95 was used, and default values were used for all other parameters. Fifty thousand cells of Panc 10.05, PATU-8988S, and PATU-8988T cell lines were harvested to construct ATAC-Seq libraries ^28^. The libraries were sequenced using Illumina NovaSeq. About 60 million reads were generated from each library. The paired-end reads were aligned to the reference genome by hisat2. Hi-C and ATAC-Seq read coverages were generated by deepTools with 10 bp bin-size, RPGC normalization, and an effective genome size of 2,864,785,220.

### Cell lines

HEK293T, PANC-1, and PATU-8988T cells were obtained from Dr. Alexander Muir (University of Chicago). Panc 10.05 was purchased from ATCC (American Type Culture Collection, USA) (https://www.atcc.org/products/crl-2547) and PATU-8988S was purchased from DSMZ (https://www.dsmz.de/collection/catalogue/details/culture/ACC-204). All cell lines were cultured at 37°C/5% CO_2_. HEK293T cells and PANC-1 cells were cultured in Dulbecco’s Modified Eagle Medium (DMEM) (Gibco, 21041025) containing 10% fetal bovine serum (FBS) (Gibco, A4766), and Panc 10.05 cells were cultured in RPMI-1640 medium (Gibco, 11875093) containing 10% FBS, as per ATCC instructions (https://www.atcc.org/products/crl-3216, https://www.atcc.org/products/crl-1469, https://www.atcc.org/products/crl-2547). PATU-8988T and PATU-8988S cells were cultured with DMEM containing 5% FBS, 5% horse serum (Gibco, 26050088), and 2 mM L-glutamine as recommended by DSMZ (Deutsche Sammlung von Mikroorganismen and Zellkulturen, Germany) (https://www.dsmz.de/collection/catalogue/details/culture/ACC-162). All cell lines have been regularly monitored and tested negative for mycoplasma using a mycoplasma detection kit (Lonza, LT07-218).

#### *TOB1-AS1* and luciferase overexpression

A 1,351 bp *TOB1-AS1* complementary DNA (cDNA) (ENST00000416263.3) was synthesized by GenScript (New Jersey, USA) and subcloned into the lentiviral pCDH-CMV-MCS-EF1-Puro plasmid (SBI, CD510B-1). The cDNA sequence in the plasmid was verified by Sanger sequencing at University of Chicago Medicine Comprehensive Cancer Center core facility. The *TOB1-AS1* overexpression plasmid was amplified by transforming Stellar™ Competent Cells (Takara, 636763) with the plasmid as per instructions and isolated by QIAGEN HiSpeed Plasmid Midi Kit (QIAGEN, 12643). LucOS-Blast vector was obtained from Dr. Yuxuan Phoenix Miao (University of Chicago), cloned, and amplified as described above.

HEK293T cells were plated in T-25 flasks and grown to 75% confluence prior to transfection. For each T-25 flask, 240μl Opti-MEM (Gibco, 31985070), 1.6μg pCMV-VSV-G, 2.56μg pMDLg/pRRE, 2.56μg pRSV-Rev, 3.4μg *TOB1-AS1* overexpression vector and 22.8μl TransIT-LT1 Transfection Reagent (Mirus, MIR 2306) were mixed and incubated at room temperature for 30 minutes, then added to the plated HEK293T cells with fresh medium. The luciferase vector was packaged into lentivirus with the same method. Upon 48 hours of incubation, lentiviral supernatant was collected, filtered through 0.45-μmpolyvinylidene difluoride filter (Millipore), and mixed with 8μg/ml polybrene. PANC-1 or PATU-8988T cells at 60% confluence were transduced with the lentiviral supernatant for 48 hours followed by three rounds of antibiotic selection with 4μg/ml puromycin for *TOB1-AS1* overexpression and 10μg/ml blasticidin for the luciferase expression. *TOB1-AS1* expression was validated by quantitative reverse transcription polymerase chain reaction (qRT-PCR), and luciferase expression was validated by in vitro bioluminescence imaging in black wall 96-well plates (Corning, 3603). D-luciferin potassium salt (Goldbio, LUCK-100) solution with 0, 1.25, 2.5, 5 and 10μl 15mg/ml was added into the wells as serial dilutions, and imaging was obtained after 5 minutes. Finally, *TOB1-AS1* overexpression or empty pCDH transduced cell lines with luciferase co-expression were built for both PATU-8988T and PANC-1 cells.

#### *TOB1-AS1* transient knock-down using antisense oligonucleotides (ASOs)

Three Affinity Plus® ASOs were synthesized by Integrated DNA Technologies (IDT), with two targeting *TOB1-AS1* and one non-targeting negative control. The ASO sequences were:

Non-targeting ASO (NC): 5’- GGCTACTACGCCGTCA- 3’

*TOB1-AS1* ASO1: 5’- GCCGATTTGGTAGCTA- 3’

*TOB1-AS1* ASO2: 5’- CTGCGGTTTAACTTCC- 3’

The ASOs were transfected into PATU-8988S and Panc 10.05 cells with Lipofecatmine^TM^ 2000 (Invitrogen, 11668019) using reverse-transfection method according to IDT protocol (https://www.idtdna.com/pages/products/functional-genomics/antisense-oligos) with a final ASO concentration of 9 nM. Cells were transfected in 6-well plates and incubated for 48 hours to reach 60% confluence before RNA extraction or Transwell assay.

### RNA isolation and qRT-PCR

Cells were plated in 6-well plates and allowed to reach 80% confluence, or transfected by ASOs as described above, prior to RNA extraction. After cells lysis in 300μl/well TRYzol^TM^ (Invitrogen, 15596026), RNA samples were prepared following the Direct-zol RNA Miniprep kit manual (RPI, ZR2052). Reverse transcription was performed using Applied Biosystems High-Capacity cDNA Reverse Transcription Kit (43-688-14) following manufacturer’s instructions. Quantitative PCR (qPCR) was conducted on StepOnePlus Real-Time PCR System (Applied Biosystems, 4376600), using PowerUp SYBR Green Master Mix (A25742) following the manufacturer’s instructions with a primer concentration of 300nM in 10μl reaction systems. Primers were ordered from Integrated DNA Technologies. Primer sequences used in this study are as follows:

*TOB1* forward: 5’-GGCACTGGTATCCTG AAA AGCC-3’

*TOB1* reverse: 5’ – GTGGCAGATTGCCACGAACATC-3’

*TOB1-AS1* forward: 5’-GGAGTGGTCAGGTGACTGATT-3’

*TOB1-AS1* reverse: 5’-ATTCCACTCCTGTTTGCAACT-3’

*GAPDH* forward: 5’– ACCACAGTCCATGCCATCAC-3’

*GAPDH* reverse: 5’-TCCACCACCCTGTTGCTGTA-3’

Relative expression levels for *TOB1-AS1* and *TOB1* were calculated by the 2^(-ΔΔC_T_) method based on *GAPDH* expression as an endogenous control.

### Transwell assay for cell invasion in vitro

Transparent PET membrane culture inserts of 24-well plate (Falcon, 353097) were coated with Cultrex Reduced Growth Factor Basement Membrane Extract (BME) (R&D Systems, 3533-010-02) at 50μg per membrane (200μl of 0.25mg/ml BME stock per membrane) at 37°C for an hour. A total of 100,000 PANC-1 cells/well, 50,000 PATU-8988T cells/well, 50,000 Panc 10.05 cells/well, or 50,000 PATU-8988S cells were resuspended in serum-free, phenol-red free DMEM medium and seeded into the coated inserts. Phenol-red free DMEM of 500μl (Gibco, A1443001) with 10% FBS was added to the bottom of the wells and the cells were allowed to invade for 16 hours. Additional wells with 500μl serum-free, phenol-red free DMEM medium without FBS in the bottom chamber were seeded with the same number of cells as indicated above as a negative control. At the end of the assay, the membranes were stained with 500μl 4μg/ml Calcein AM (CaAM) (Corning, 354216) for one hour at 37°C. The cells that failed to invade were removed from the top chamber with a cotton swab and all inserts were transferred into 1x Cell Dissociation Solution (Bio-Techne, 3455-05-03) and shaken at 150rpm for an hour at 37°C. Finally, CaAM signal from the invaded cells was measured by a plate reader (Perkin Elmer Victor X3) at 465/535nm.

### Tumor metastasis in vivo

All animal experiments for this study were approved by the University of Chicago Institutional Animal Care and Use Committee (IACUC) prior to execution. Male NOD *scid* gamma (NSG) mice were ordered from the Jackson Laboratory (strain#005557). For tail vein inoculation, mice were injected intravenously through the tail vein with luciferase-expressing at 400,000 cells/mouse for PANC-1 cells in cold phosphate buffered saline (PBS) (Gibco, 10010-023). For orthotopic inoculation, mice were injected with 200,000 PANC-1 cells/mouse into the pancreas under general anesthesia. Cells were resuspended in cold PBS containing 5.6mg/mL Cultrex Reduced Growth Factor BME (R&D Systems, 3533-010-02). Primary tumor and metastatic tumor burdens were measured weekly for 4 and 6 weeks for tail vein injection models and orthotopic models, respectively, via bioluminescence imaging using Xenogen IVIS 200 Imaging System (PerkinElmer) at the University of Chicago Integrated Small Animal Imaging Research Resource (iSAIRR) Facility. Each mouse was weighed and injected intra-peritoneally with D-luciferin solution at a concentration of 150μg/g of body weight 14 minutes prior to image scanning ventral side up.

### Ex vivo IVIS imaging

Ex vivo imaging was done for the PANC-1 orthotopic injection mice after 8 weeks of orthotopic inoculation. Mice were injected intra-peritoneally with D-luciferin solution at a concentration of 150μg/g of body weight immediately before euthanasia. Immediately after necropsy, mice were dissected, and tissues of interest (primary tumors, livers and spleens) were placed into individual wells of 6-well plates covered with 300 μg/mL D-luciferin. Tissues were imaged using Xenogen IVIS 200 Imaging System (PerkinElmer) and analysis was performed (Living Image Software, PerkinElmer) maintaining the regions of interest (ROIs) over the tissues as a constant size.

### Tumor RNA sequencing and gene expression analysis

RNA was isolated from mouse subcutaneous tumors (six *TOB1-AS1* overexpression and six control mice) after 6 weeks of PANC-1 cell subcutaneous injection using Direct-zol RNA Miniprep kit (RPI, ZR2052). The quality and quantity of the RNA were assessed using Qubit. Sequencing was performed using the Illumina NovaSeq 6000. About 40 million reads were sequenced per sample. The pair-end reads were aligned to mouse genome (mm10) and human genome (hg19) with hisat2, and the reads mapped to mouse or human genomes were disambiguated using AstraZeneca-NGS disambiguate package. Gene counts were generated with htseq-count. Differential gene expression was analyzed using DESeq2. Differentially expressed genes were defined as genes with an FDR smaller than 0.1 and a fold change greater than 1.5.

### Code availability

The HYENA package is available at https://github.com/yanglab-computationalgenomics/HYENA.

## Results

### HYENA workflow

Conceptually, the SVs leading to elevated gene expression are eQTLs. The variants are SVs instead of commonly used germline single nucleotide polymorphisms (SNPs) in eQTL analysis. With somatic SVs and gene expression measured from the same tumors through WGS and RNA-Seq, we can identify enhancer hijacking target genes by eQTL analysis. However, the complexities of cancer and SVs pose many challenges. For instance, there is tremendous inter-tumor heterogeneity—no two tumors are identical at the molecular level. In addition, there is substantial intra-tumor heterogeneity as tumor tissues are always mixtures of tumor, stromal, and immune cells. Moreover, genome instability is a hallmark of cancer, and gene dosages are frequently altered ^29^. Furthermore, gene expression networks in cancer are widely rewired ^30^, and outliers of gene expression are common.

Here, we developed an algorithm HYENA to overcome the challenges described above (see more details in Methods Section). We used a gene-centric approach to search for elevated expression of genes correlated with the presence of SVs within 500 kb of transcription start sites (**Fig. 1B**). Although promoter-enhancer interaction may occur as far as several mega-bases, mega-base-level long-range interactions are extremely rare. In addition, although duplicated enhancers can upregulate genes ^31,32^, we do not consider these as enhancer hijacking events since no neo-promoter-enhancer interactions are established. However, small deletions can remove TAD boundaries or repressive elements and lead to neo-promoter-enhancer interactions (**Fig. 1A**). Therefore, small tandem duplications were discarded, and small deletions were retained. For each gene, we annotated SV status (presence or absence of nearby SVs) for all samples. Samples in which the testing genes were highly amplified were discarded since many of these genes are amplified by circular extrachromosomal DNA (ecDNA) ^33^, and ecDNA can promote accessible chromatin ^34^ with enhancer rewiring ^35^. Only genes with nearby SVs in at least 5% of tumors were further considered. In contrast to CESAM and PANGEA, we did not use linear regression to model the relationships between SV status and gene expression because linear regression is sensitive to outliers and many false positive associations would be detected ^36^. Instead, we used a rank-based normal-score regression approach. After quantile normalization of gene expression for both protein-coding and non-coding genes, we ranked the genes based on quantile-normalized expression and transformed the ranks to the quantiles of the standard normal distribution. We used the z scores (normal scores) of the quantiles as dependent variables in regression. In the normal-score regression model, tumor purity, copy number of the tested gene, patient age, and sex were included as covariates since these factors confound gene expression. We also included gene expression principal components (PCs) that were not correlated with SV status to model unexplained variations in gene expression. To deduce a better null distribution, we permuted the gene expression 100 to 1000 times (**Supplementary Table S1**) and ran the same regression models. All *P* values from the permutations were pooled together and used as the null distribution to calculate empirical *P* values. Then, multiple testing corrections were performed on one-sided *P* values since we are only interested in elevated gene expression under the influence of nearby SVs. Finally, genes were discarded if their elevated expression could be explained by germline eQTLs. The remaining genes were candidate enhancer hijacking target genes.

### Benchmarking performances

There is no gold standard available to comprehensively evaluate the performance of HYENA. We compared HYENA’s performance to two other algorithms—CESAM and PANGEA. All three algorithms were run on the same somatic SVs and gene expression data from six types of adult tumors profiled by the PCAWG (**Supplementary Table S1**): malignant lymphoma (MALY), stomach/gastric adenocarcinoma (STAD), chromophobe renal cell carcinoma (KICH), colorectal cancer (COAD/READ), thyroid cancer (THCA), and lung squamous cell carcinoma (LUSC) ^21^. Note that PANGEA depends on promoter-enhancer interactions predicted from cell lines, and such data were not available for thyroid tissue. Therefore, thyroid cancer data were not analyzed by PANGEA. To compare the performance of HYENA to the other algorithms, we used the following three strategies.

First, we used eight known enhancer hijacking target genes including *MYC* ^9^, *BCL2* ^8^, *CCNE1* ^37^, *TERT* ^7^, *IGF2* ^13,37^ (in two tumor types), *IGF2BP3* ^38^ and *IRS4* ^13^ to test the sensitivities. Out of the eight known enhancer hijacking genes, HYENA detected four (*MYC*, *BCL2*, *TERT*, and *IGF2BP3*) (**Fig. 2A** and **Supplementary Fig. S1A**), CESAM detected three (*MYC*, *BCL2*, and *TERT*), and PANGEA did not detect any (**Fig. 2A**). In the five tumor types analyzed by all three algorithms, HYENA identified a total of 25 candidate genes, CESAM identified 19, whereas PANGEA identified 255 genes (**Fig. 2B**, **Supplementary Tables S6**, **S7**, and **S8**). Six genes were detected by both HYENA and CESAM, while PANGEA had little overlap with the other algorithms (**Fig. 2B**). The ability of the algorithms to detect known target genes seems to be sensitive to sample size. Both *IGF2* and *IRS4* were initially discovered by CESAM as enhancer hijacking target genes using CNV breakpoints profiled by microarray with much larger sample sizes (378 colorectal cancers and 497 lung squamous cell carcinomas) ^13^. In the PCAWG, there were many fewer samples with both WGS and RNA-Seq date available (51 colorectal cancers and 47 lung squamous cell carcinomas). Neither *IGFR* nor *IRS4* was detected by any algorithms. *IGF2* reached 5% SV frequency cutoff required by HYENA, however its FDR did not reach the significance cutoff (**Supplementary Fig. S1B**). In stomach/gastric adenocarcinoma, *IGF2* and *CCNE1* were identified as enhancer hijacking target genes in a cohort of 208 samples ^37^. Neither of these genes was detected by any algorithms because there were only 29 stomach tumors in the PCAWG. Therefore, known target genes missed by HYENA were likely due to small sample size. In summary, HYENA had the best sensitivity of the three algorithms.

**Figure 2.**
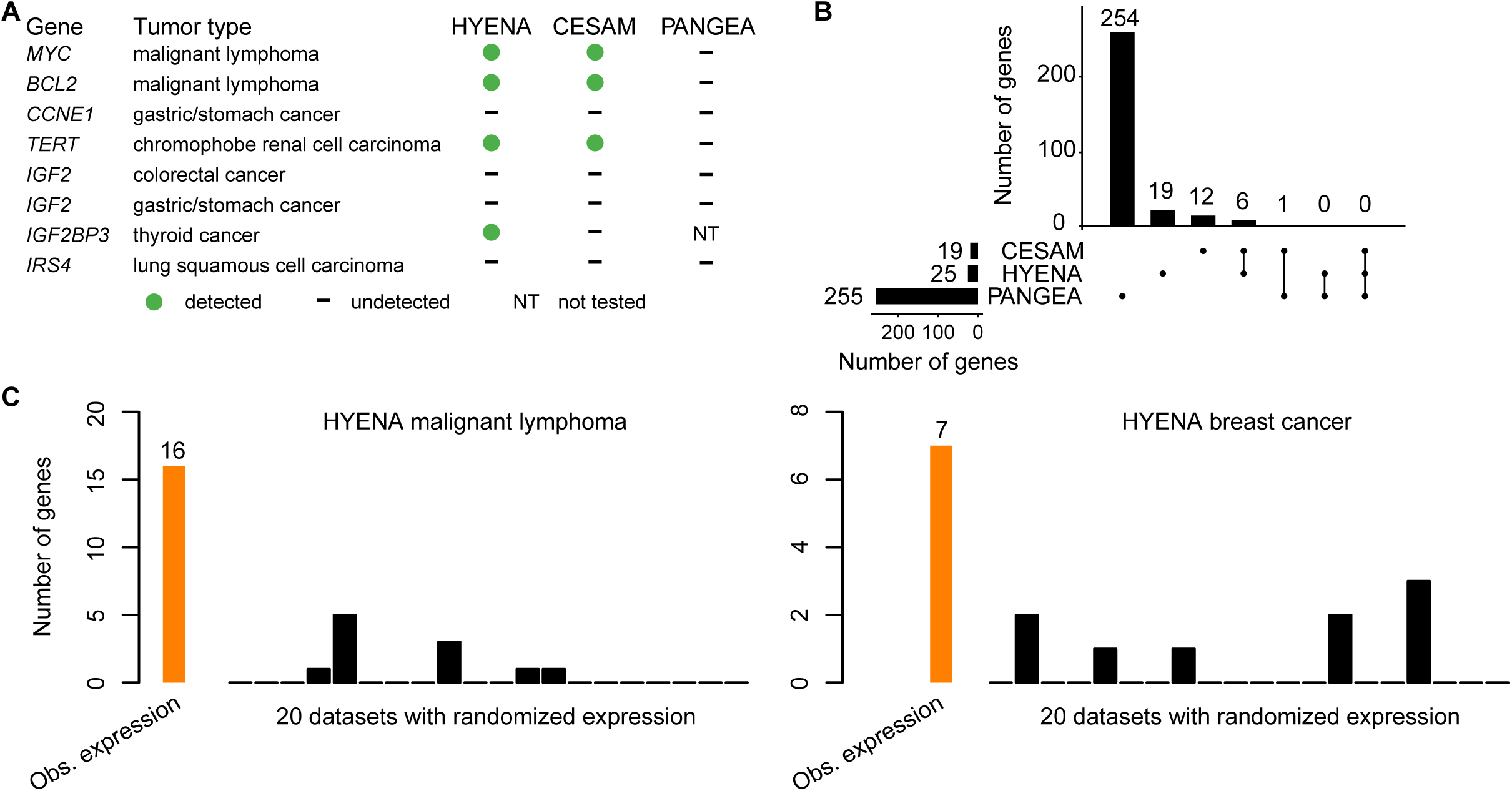
Benchmarking HYENA. **A**, Comparison of HYENA, CESAM, and PANGEA in detecting oncogenes known to be activated by enhancer hijacking in six tumor types from the PCAWG cohort. **B**, UPSET plot demonstrating candidate genes identified and shared among the three tools in five tumor types of PCAWG. The numbers of candidate genes predicted by three algorithms are shown on the bottom left (19, 25, and 255). On the bottom right, individual dots denote genes detected by one tool, and dots connected by lines denote genes detected by multiple tools. The numbers of genes detected are shown above the dots and lines. For example, the dot immediately on the right of “PANGEA” shows there are 254 candidate genes detected only by PANGEA but not CESAM and HYENA. The left most line connecting two dots indicates that there are six genes detected by both CESAM and HYENA but not by PANGEA. **C**, Number of genes detected by HYENA in two PCAWG tumor types using observed gene expression and randomized expression. Genes detected in random expression datasets are false positives.

Second, we also expect immunoglobulin genes to be detected as enhancer hijacking candidates in B-cell lymphoma due to V(D)J recombination. In B cells, V(D)J recombination occurs to join different variable (V), joining (J), and constant (C) segments to produce antibodies with a wide range of antigen recognition ability. Therefore, certain segments have elevated expression and the recombination events can be detected as somatic SVs. Of the 16 genes detected by HYENA in malignant lymphoma (B-cell derived Burkitt lymphomas ^39^), there were two immunoglobulin light chain genes from the lambda cluster (*IGLC7* and *IGLJ7*) and an immunoglobulin-like gene *IGSF3* (**Supplementary Table S6**). CESAM detected 11 genes, one of which was an immunoglobulin gene (*IGLC7*) (**Supplementary Table S7**). In contrast, PANGEA detected 30 candidate genes, but none were immunoglobulin genes (**Supplementary Table S8**). These data further support HYENA as the algorithm with the best sensitivity among the three algorithms.

Third, to evaluate the specificity of the algorithms, we ran each algorithm on 20 datasets generated by randomly shuffling gene expression data in both MALY and breast cancer (BRCA). Since these gene expression data were random, there should be no associations between SVs and gene expression, and all genes detected should be false positives. In malignant lymphoma with observed gene expression, HYENA, CESAM, and PANGEA detected 16, 11, and 30 candidate genes respectively (**Supplementary Tables S6, S7 and S8**). In the 20 random gene expression datasets for malignant lymphoma, HYENA detected an average of 0.55 genes per dataset (**Fig. 2C**), and CESAM detected an average of 0.5 genes per dataset, whereas PANGEA detected an average of 40 genes per dataset (**Supplementary Fig. S2**). In breast cancer with observed gene expression, HYENA, CESAM, and PANGEA detected 7, 9, and 2,309 candidate genes, respectively (**Supplementary Tables S6, S7 and S8**). In 20 random gene expression datasets for breast cancer, HYENA, CESAM, and PANGEA detected 0.45, 0.9 and 2,296 genes on average (**Fig. 2C** and **Supplementary Fig. S2**). In both tumor types, the numbers of false positives called by PANGEA in random datasets were comparable to the numbers of genes detected with observed gene expression (**Supplementary Fig. S2**). In summary, HYENA predicted the least number of false positives among the three algorithms.

Overall, HYENA has superior sensitivity and specificity in the detection of enhancer hijacking genes. Although the performances of CESAM were similar to HYENA, the genes detected by HYENA and CESAM in the six benchmarking tumor types had little overlap (**Fig. 2B**). We performed extensive validation on one gene detected only by HYENA.

### Enhancer hijacking candidate genes in the PCAWG

We used HYENA to analyze a total of 1,146 tumors across 25 tumor types in the PCAWG with both WGS and RNA-Seq data. When each tumor type was analyzed individually, we identified 108 candidate enhancer hijacking target genes in total (**Supplementary Tables S1** and **S6**), four of which were known enhancer hijacking targets (**Fig. 3A**). *TERT* was detected in kidney cancers both from the US cohort (KICH) and the European cohort (RECA) which further demonstrated the reproducibility of HYENA. All other candidate genes were only detected in one tumor type, highlighting high tumor type specificity of the findings. The number of genes detected in each tumor type also differed dramatically (**Fig. 3B**). No genes were detected in bladder cancer (BLCA), cervical cancer (CESC), glioblastoma multiforme (GBM), or low-grade glioma (LGG), probably due to their small sample sizes. Pancreatic cancer (PACA) had the greatest number of candidate genes. There were two liver cancer cohorts with comparable sample sizes—LIHC from the US and LIRI from Japan. Interestingly, a total of 14 genes were identified in the US cohort whereas no genes were found in the Japanese cohort. One possible reason for such a drastic difference could be that hepatitis B virus (HBV) infection is more common in liver cancer in Japan ^40^, and virus integration into the tumor genome can result in oncogene activation ^41^. In Chronic Lymphocytic Leukemia (CLLE), a total of six genes were detected, and three were immunoglobulin genes from both the lambda and kappa clusters (**Supplementary Table S6**). Given that sample size and genome instability can only explain a small fraction of the variations of enhancer hijacking target genes detected in different tumor types, the landscape of enhancer hijacking in cancer seems to be mainly driven by the underlying disease biology. Intriguingly, out of the 108 candidate genes, 54 (50%) were non-coding genes including lncRNAs and microRNAs (**Fig. 3B**).

**Figure 3.**
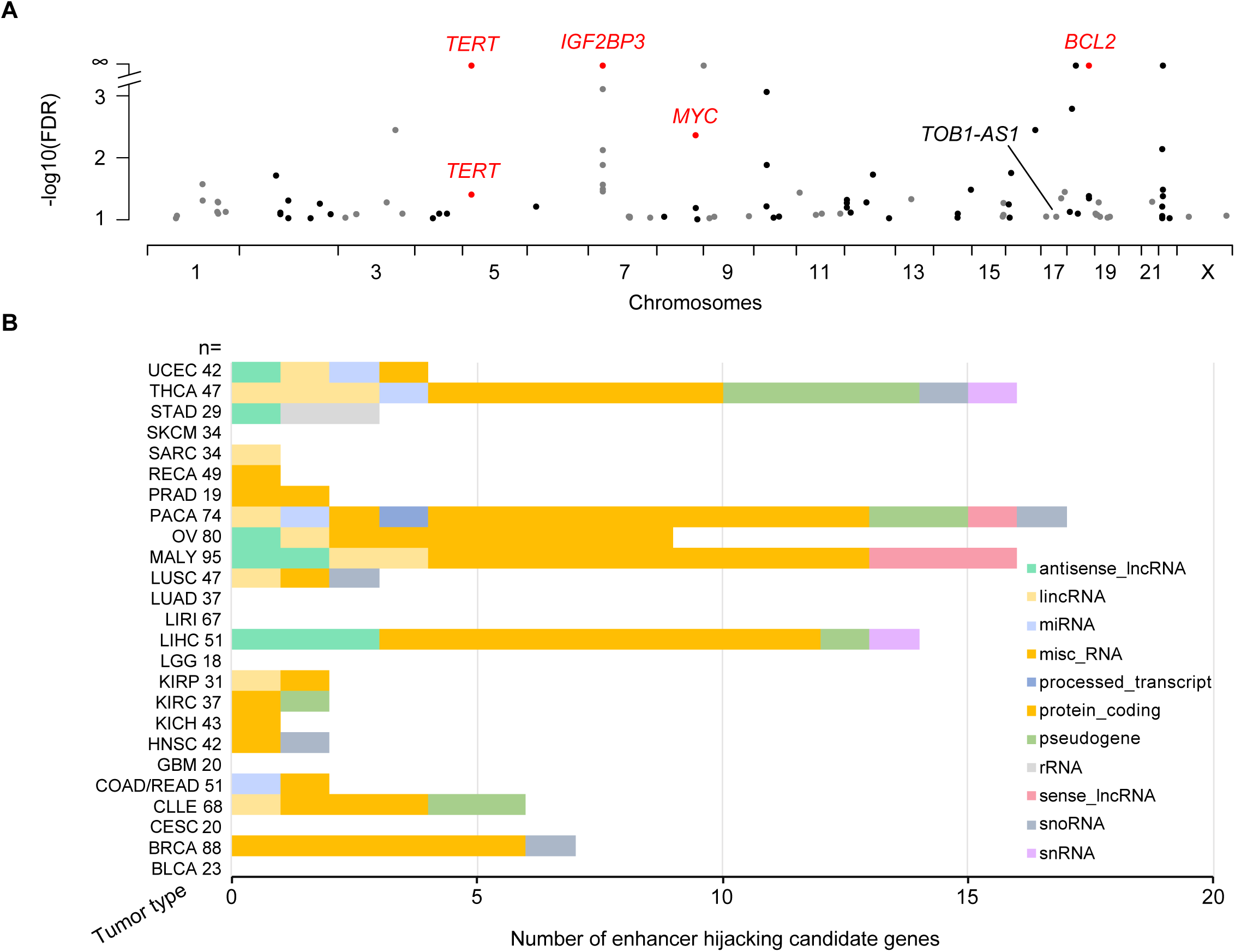
Enhancer hijacking candidate genes in PCAWG. **A**, Candidate genes detected by HYENA in individual tumor types of PCAWG. *TERT* is plotted twice since it is detected in two cancer types. Genes labelled as red are known enhancer hijacking targets. **B**, Diverse types of candidate genes identified by HYENA in PCAWG. Numbers after tumor type names denote sample size in the corresponding tumor types.

### Neo-TADs formed through somatic SVs

Next, we focused on the most frequently altered candidate non-coding enhancer-hijacking target gene in pancreatic cancer: *TOB1-AS1* (**Fig. 4A**), a lncRNA. *TOB1-AS1* was not detected as a candidate gene by either CESAM (**Supplementary Table S7**) or PANGEA (**Supplementary Table S8**) using the same input data. Seven (9.6%) out of 74 tumors had some form of somatic SVs near *TOB1-AS1* including translocations, deletions, inversions, and tandem duplications (**Fig. 4B** and **Supplementary Table S9**). For example, tumor 9ebac79d-8b38-4469-837e-b834725fe6d5 had a translocation between chromosomes 17 and 19 (**Fig. 4C**). The breakpoints were upstream of *TOB1-AS1* and upstream of *UQCRFS1* (**Fig. 4D**). In tumor 748d3ff3-8699-4519-8e0f-26b6a0581bff, there was a 19.3 Mb deletion which brought *TOB1-AS1* next to a region downstream of *KCNJ2* (**Fig. 4C** and **4E**).

**Figure 4.**
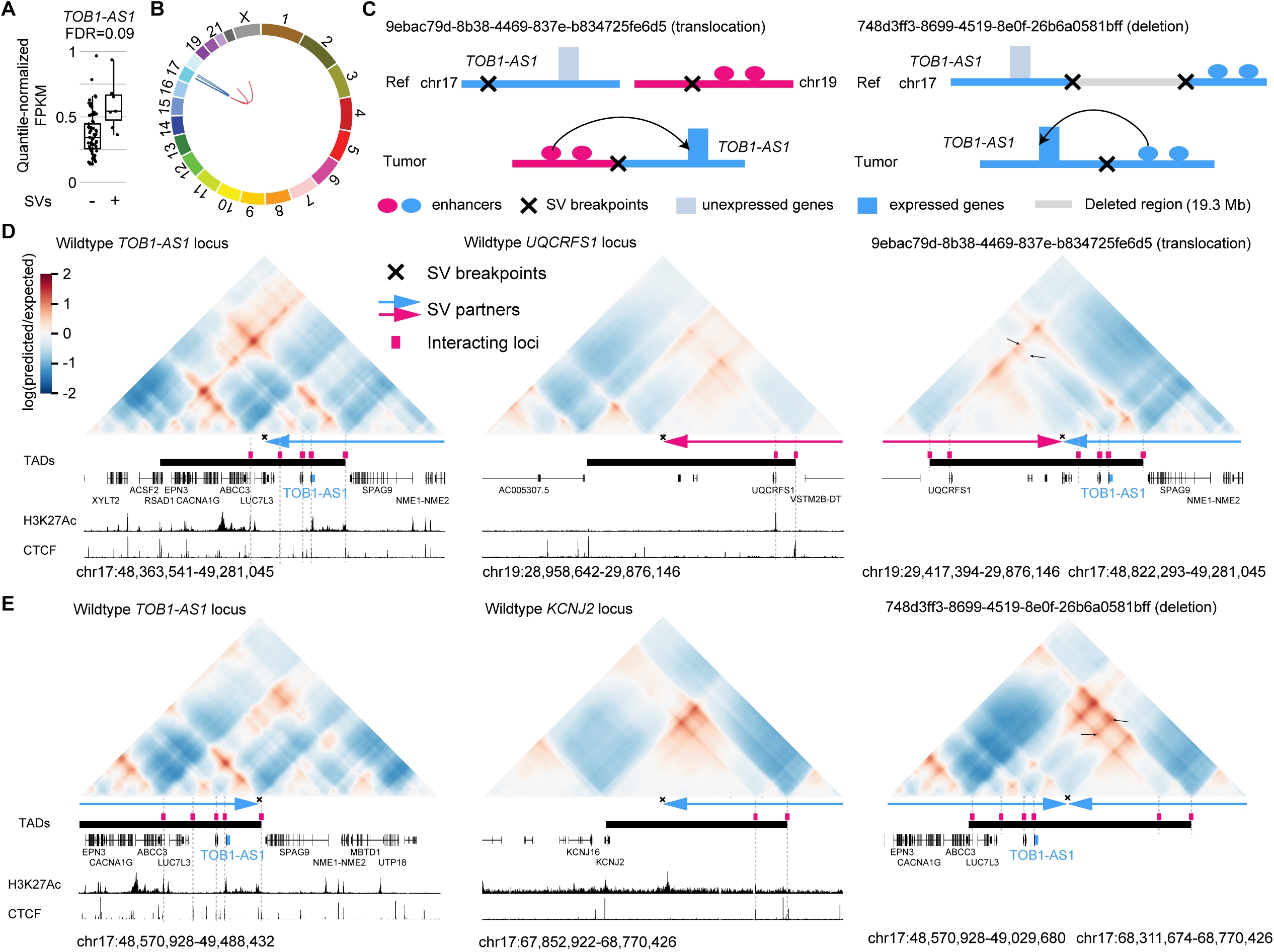
*TOB1-AS1* activated by various types of SVs in pancreatic cancer. **A**, Normalized expression of *TOB1-AS1* in samples with (n=7) and without (n=67) nearby SVs in pancreatic cancers. The boxplot shows median values (thick black lines), upper and lower quartiles (boxes), and 1.5× interquartile range (whiskers). Individual tumors are shown as black dots. **B**, Circos plot summarizing intrachromosomal SVs (blue, n=5) and translocations (red, n=3) near *TOB1-AS1*. **C**, Diagrams depicting putative enhancer hijacking mechanisms that activate *TOB1-AS1* in one tumor with a 17:19 translocation (left panel) and another tumor with a large deletion (right panel). **D**, Predicted 3D chromatin interaction maps of *TOB1-AS1* (left panel), *UQCRFS1* (middle panel), and the translocated region in tumor 9ebac79d-8b38-4469-837e-b834725fe6d5 (right panel). The downstream fragment of the chromosome 19 SV breakpoint was flipped in orientation and linked to chromosome 17. H3K27Ac and CTCF ChIP-Seq data of PANC-1 cell line are shown at the bottom. The expected level of 3D contacts depends on linear distance between two genomic locations. Longer distances correlate with fewer contacts. Akita predicts 3D contacts based on DNA sequences. The heatmaps are showing the ratio between predicted and expected contacts. The darkest red represent regions having 100 times more contacts than expected given the distance between the regions. **E**, Predicted 3D chromatin interaction maps of *TOB1-AS1* (left panel) and *KCNJ2* (middle panel) loci without deletion as well as the same region following deletion in tumor 748d3ff3-8699-4519-8e0f-26b6a0581bff (right panel).

We used Akita ^23^, a convolutional neural network that predicts 3D genome organization, to assess the 3D architecture of the loci impacted by SVs. While 3D structures are dynamic and may change with cell-type and gene activity, TAD boundaries are often more stable and remain similar across different cell-types ^1^. TAD boundaries are defined locally by the presence of binding sites for CTCF, a ubiquitously expressed DNA-binding protein ^1,26^, and TAD formation arises from the stalling of the cohesin-extruded chromatin loop by DNA-bound CTCF at these positions ^42^. For this reason, it is expected that upon chromosomal rearrangements, normal TADs can be disrupted, and new TADs can form by relocation of TAD boundaries. This assumption has been validated with direct experimental evidence from examining the “neo-TADs” associated with SVs at different loci ^43–45^. The wildtype *TOB1-AS1* locus had a TAD between a CTCF binding site in *RSAD1* and another one upstream of *SPAG9* (**Fig. 4D** and **Supplementary Fig. S3**). There were TADs spanning *UQCRFS1* and downstream of *KCNJ2* in the two partner regions (**Fig. 4D**, **4E** and **Supplementary Fig. S3**). In tumor 9ebac79d-8b38-4469-837e-b834725fe6d5, the translocation was predicted to lead to a neo-TAD resulting from merging the TADs of *TOB1-AS1* and *UQCRFS1* (**Fig. 4D)**. In tumor 748d3ff3-8699-4519-8e0f-26b6a0581bff, another neo-TAD was predicted to form as a result of the deletion that merged the TADs of *TOB1-AS1* and the downstream portion of *KCNJ2* (**Fig. 4E**). In both cases, within these predicted neo-TADs, Akita predicted strong chromatin interactions involving several CTCF binding sites and H3K27Ac peaks between *TOB1-AS1* and its two SV partners (**Fig. 4D** and **4E** black arrows in the right panels), indicating newly formed promoter-enhancer interactions. In the vicinity of the *TOB1-AS1* locus, *TOB1-AS1* was the only gene with significant changes in gene expression. Similar neo-TADs could be observed in two additional tumors (**Supplementary Fig. S4**). In two tumors harboring tandem duplications of *TOB1-AS1* of 317 kb and 226 kb, the *TOB1-AS1* TADs were expanded (**Supplementary Fig. S5A**). However, not all SVs near *TOB1-AS1* led to alterations in TAD architecture; for example, in tumor a3edc9cc-f54a-4459-a5d0-097879c811e5, *TOB1-AS1* was predicted to remain in its original TAD after a 4 Mb tandem duplication (**Supplementary Fig. S5B**). In summary, at least four out of the seven tumors harboring somatic SVs near *TOB1-AS1* were predicted to result in neo-TADs including *TOB1-AS1*. We then used another deep-learning algorithm called Orca ^25^ to predict 3D genome structure based on DNA sequences. Orca-predicted 3D genome architectures were very similar to Akita predictions (**Supplementary Fig. S6**) in neo-TAD formation due to SVs in the *TOB1-AS1* locus.

To further study the 3D genome structure of the *TOB1-AS1* locus, we performed high-resolution in situ Hi-C sequencing for four pancreatic cancer cell lines. Among these, two cell lines (Panc 10.05 and PATU-8988S) had high expression of *TOB1-AS1*, whereas the other two (PANC-1 and PATU-8988T) had low expression (**Fig. 5A**). At mega-base-pair scale, three cell lines (Panc 10.05, PATU-8988S, and PATU-8988T) carried several SVs (black arrows in **Fig. 5B**). In Panc 10.05, a tandem duplication (chr17:43,145,000-45,950,000) was observed upstream of *TOB1-AS1* (**Fig. 5B** black arrow in the left most panel and **Supplementary Table S10**). However, the breakpoint was too far away (2 Mb) from *TOB1-AS1* (chr17:48,944,040-48,945,732) and unlikely to regulate its expression. A neo chromatin loop was detected by NeoLoopFinder ^20^ near *TOB1-AS1* (chr17:34,010,000-48,980,000) driven by a deletion (chr17:34,460,000-47,450,000) detected by EagleC ^27^ (**Supplementary Fig. S7A**, **Supplementary Tables S5** and **S10**). The deletion breakpoint was also too far away (1.5 Mb) from *TOB1-AS1* and unlikely to regulate its expression. No other SVs or neo chromatin loops were detected near *TOB1-AS1* (**Supplementary Tables S5** and **S10**). Interestingly, there was a CNV breakpoint (chr17:48,980,000) 36 kb downstream of *TOB1-AS1* in Panc 10.05 (**Fig. 5C** left most panel) which was also the boundary of the neo chromatin loop. In the high copy region (upstream of the CNV breakpoint), heterozygous SNPs were present with allele ratios of approximately 4:1 (**Supplementary Fig. S8A**), whereas in the low copy region (downstream of the CNV breakpoint), all SNPs were homozygous (**Supplementary Fig. S8B**). These data suggested that the DNA copy number changed from five copies to one copy at the CNV breakpoint. The gained copies must connect to some DNA sequences since there should not be any free DNA ends other than telomeres. Given that no off-diagonal 3D genome interactions were observed at chr17:48,980,000, we considered the possibilities that the high copy region was connected to repetitive sequences or to sequences that were not present in the reference genome. If so, reads mapped to the high copy region should have an excessive amount of non-uniquely mapped mates or unmapped mates. However, this was not the case (**Supplementary Fig. S9**). The only possible configuration was a foldback inversion in which two identical DNA fragments from the copy gain region were connected head to tail (**Fig. 5D** bottom left panel). As a result, in Panc 10.05, there was a wildtype chromosome 17, two foldback-inversion-derived chromosomes, and a translocation-derived chromosome (**Fig. 5D** bottom left panel and **Supplementary Fig. S7B**). Foldback inversions are very common in cancer. If DNA double strand breaks are not immediately repaired, following replication, the two broken ends of sister chromatids can self-ligate head to tail and sometimes result in dicentric chromosomes ^46,47^. Algorithms, such as hic-breakfinder ^48^ and EagleC ^27^, rely on off-diagonal 3D genomic interactions in Hi-C contact matrix to detect SVs. However, foldback inversions do not form any off-diagonal interactions since the two connected DNA fragments have the same coordinates, so they are not detectable by existing algorithms. The 3D genome structure of the *TOB1-AS1* locus in Panc 10.05 was quite distinct from the other three cell lines (**Fig. 5C**). The region immediately involved in the foldback inversion had homogeneous 3D interactions (**Fig. 5C** dashed blue triangle in the left most panel) suggesting that a neo-subdomain was formed (**Fig. 5D** right panel). The high expression of *TOB1-AS1* in Panc 10.05 was likely a combined effect of the copy gain and the neo-subdomain. In PATU-8988S and PATU-8988T, a shared SV (chr17:48,880,000-52,520,000) near *TOB1-AS1* was detected (**Fig. 5B** two right panels) since the two cell lines were derived from the same pancreatic cancer patient ^49^. This shared SV could not regulate *TOB1-AS1* because it pointed away from *TOB1-AS1* (**Supplementary Fig. S10**). No other SVs were found near *TOB1-AS1* in these two cell lines. The high expression of *TOB1-AS1* in PATU-8988S was likely due to transcriptional regulation since the promoter of *TOB1-AS1* in PATU-8988S was more accessible than that in PATU-8988T (**Fig. 5E**). This result was consistent with a handful of patient tumors that had high expression of *TOB1-AS1* without any SVs (**Fig. 4A**).

Taken together, our results demonstrated that HYENA can detect genes activated by reorganization of 3D genome architecture.

**Figure 5.**
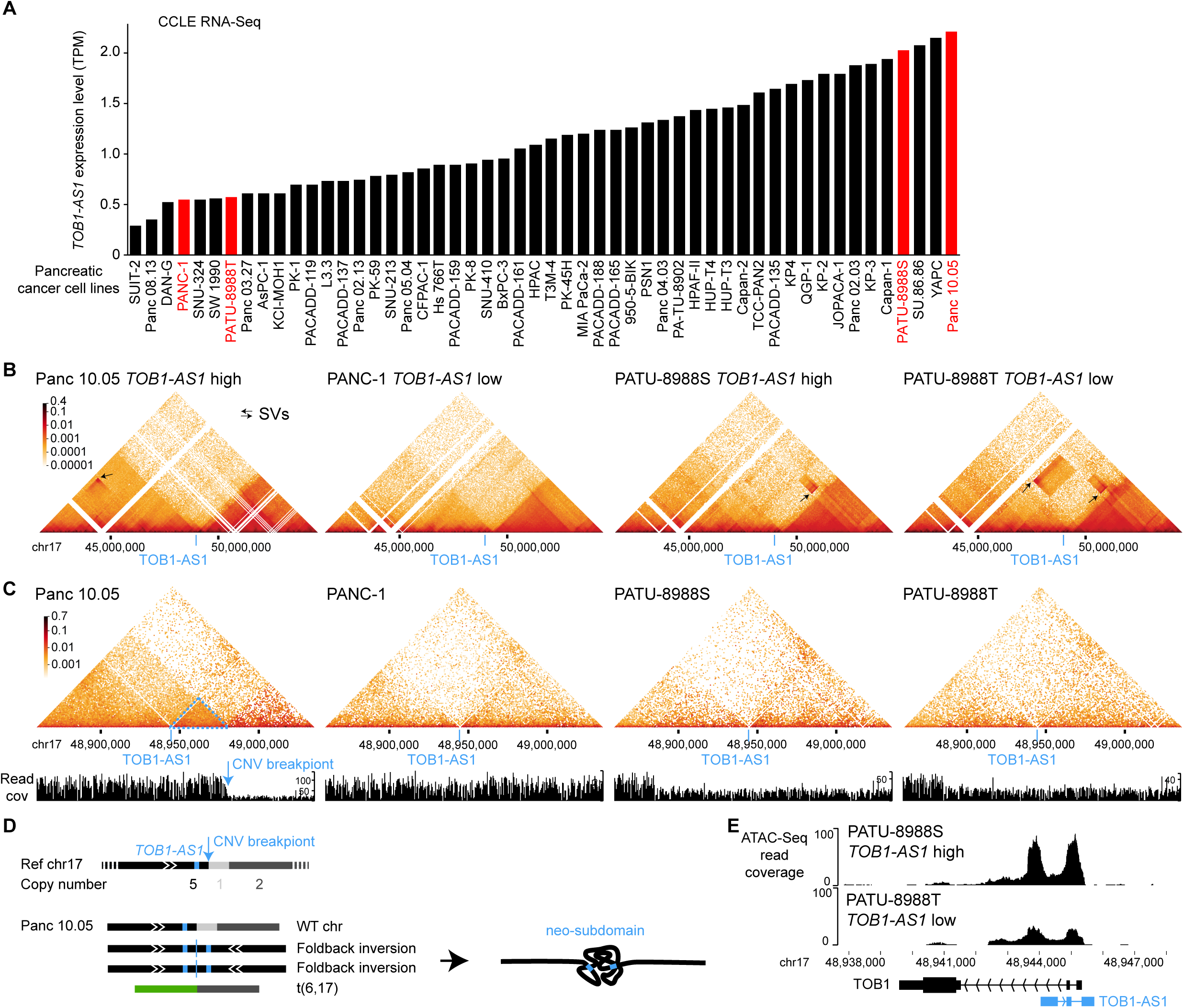
3D genome structures in the *TOB1-AS1* locus in pancreatic cancer cell lines. **A**, *TOB1-AS1* expression in pancreatic cancer cell lines in CCLE. The cell lines in red are selected for further studies. **B** and **C**, 3D genomic interactions in four pancreatic cancer cell lines. Black arrows represent SVs with off-diagonal interactions. The locations of *TOB1-AS1* are marked by blue lines. In Panc 10.05, the blue arrow points to the CNV breakpoint and the dashed blue triangle represents the neo-subdomain formed due to the foldback inversion. **D**, The reference chromosome 17 and derived chromosomes in Panc 10.05. The chromosomes are not to scale. *TOB1-AS1* is shown as small blue boxes in the chromosomes. **E**, Open chromatin measured by ATAC-Seq in PATU-8988S and PATU-8988T at the *TOB1-AS1* locus.

### Oncogenic functions of *TOB1-AS1*

*TOB1-AS1* has been reported as a tumor suppressor in several tumor types ^50,51^. However, HYENA predicted it to be an oncogene in pancreatic cancers. To test the potential oncogenic functions of *TOB1-AS1* in pancreatic cancer, we performed both in vitro and in vivo experiments. We surveyed pancreatic cancer cell line RNA-Seq data from Cancer Cell Line Encyclopedia (CCLE) and identified that the commonly transcribed isoform of *TOB1-AS1* in pancreatic cancers was ENST00000416263.3 (**Supplementary Fig. S11**). The synthesized *TOB1-AS1* cDNA was cloned and overexpressed in two pancreatic cancer cell lines, PANC-1 and PATU-8988T, both of which had low expression of *TOB1-AS1* (**Fig. 5A** and **Supplementary Fig. S12A**). In both cell lines, overexpression of *TOB1-AS1* (**Fig. 6A**) promoted in vitro cell invasion (**Fig. 6B**). In addition, three weeks after tail vein injection, PANC-1 cells with *TOB1-AS1* overexpression caused higher metastatic burden in immunodeficient mice than the control cells (**Fig. 6C**). Six weeks after orthotopic injection, mice carrying *TOB1-AS1* overexpressing PANC-1 cells showed exacerbated overall tumor burden (**Fig. 6D**), elevated primary tumor burden, and elevated metastatic burden in the spleen (**Fig. 6E** and **Supplementary Fig. S12B**). Liver metastasis was not affected (**Supplementary Fig. S12C**). In addition, we knocked down *TOB1-AS1* in two other pancreatic cancer cell lines Panc 10.05 and PATU-8988S, both of which had high expression of *TOB1-AS1* (**Fig. 5A** and **Supplementary Fig. S12A**), using two antisense oligonucleotides (ASOs) (**Fig. 6F**). *TOB1-AS1* expression was reduced by approximately 50% by both ASOs (**Fig. 6G**). Knockdown of *TOB1-AS1* substantially suppressed cell invasion in vitro (**Fig. 6H**). Note that PATU-8988T and PATU-8988S were derived from the same liver metastasis of a pancreatic cancer patient, and they had drastic differences in *TOB1-AS1* expression (**Fig. 5A** and **Supplementary Fig. S12A**). It was reported that PATU-8988S can form lung metastases in vivo with tail vein injection of nude mice, whereas PATU-8988T cannot form any metastases in any organ ^49^. By altering the expression of *TOB1-AS1*, we were able to reverse the cell invasion phenotypes in these two cell lines (**Fig. 6B** and **6H**). These results suggested that *TOB1-AS1* has an important function in regulating cell invasion.

**Figure 6.**
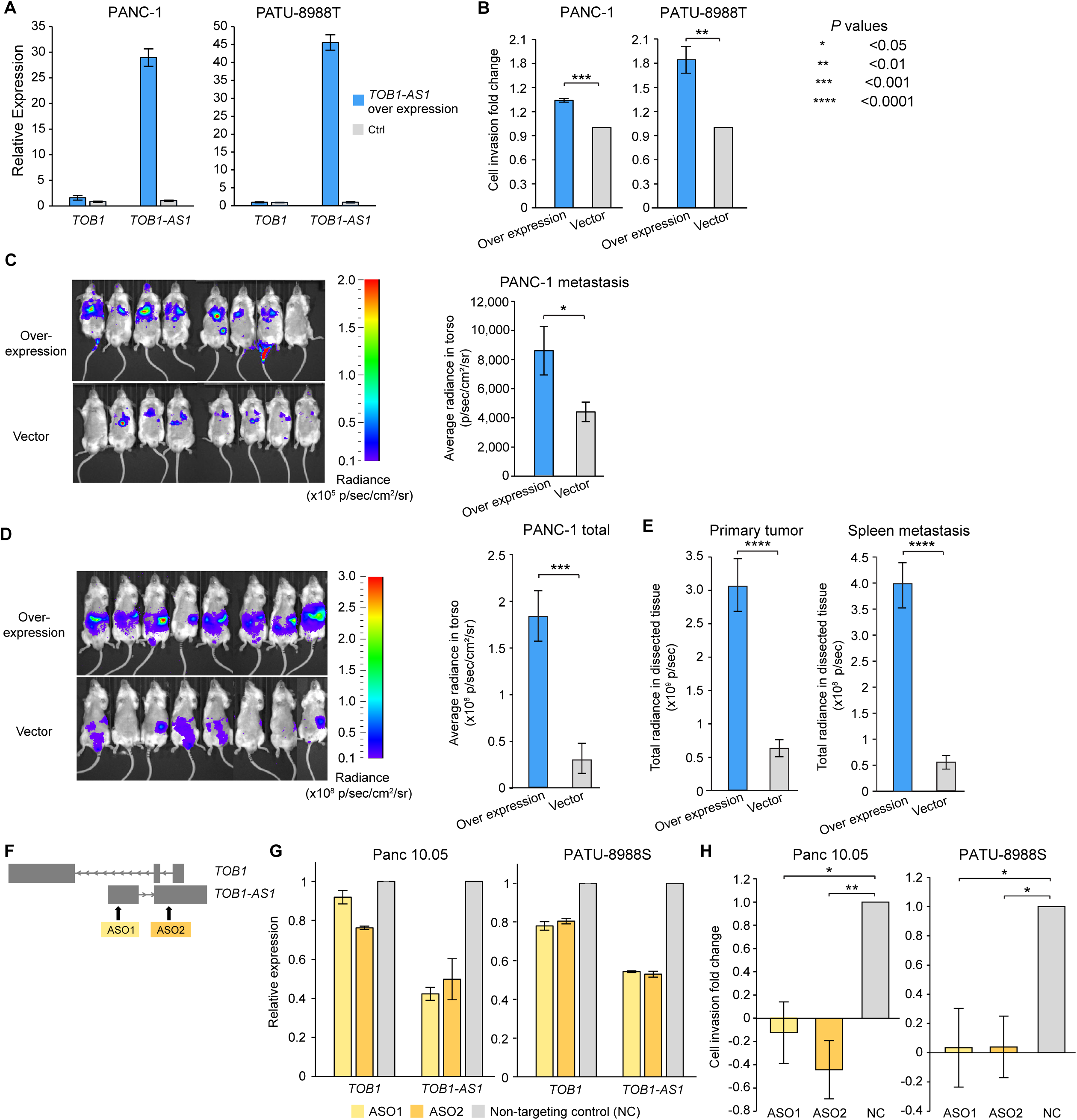
*TOB1-AS1* promotes cell invasion and tumor metastasis. **A**, *TOB1-AS1* and *TOB1* relative expression levels in PATU-8988T and PANC-1 cells transduced with *TOB1-AS1* overexpression vector (n=3) or control vector (n=3). **B**, *TOB1-AS1* overexpression in PATU-8988T (4 biological replicates) and PANC-1 (3 biological replicates) promoted in vitro cell invasion using Transwell assay. Each biological replicate was an independent experiment with 7 technical replicates per experimental group. The average fold change of cell invasion was calculated after the background invasion measured in the absence of any chemotactic agent was subtracted from each technical replicate. *P* values were calculated by two-sided student t test. **C**, *TOB1-AS1* overexpression in PANC-1 cells promoted in vivo tumor metastasis in the tail vein injection model. **D**, *TOB1-AS1* overexpression in PANC-1 cells exacerbated in vivo tumor growth and spontaneous metastasis in the orthotopic tumor model. Images of radiance in immunodeficient mice are shown on the left while the quantifications of radiance are shown on the right. Eight mice were used in both the overexpression group and the empty vector control. The images were analyzed by setting the regions of interest (ROIs) to mouse torsos and measuring the average radiance level (in p/sec/cm^2^/sr). **E**, Primary tumor burden and spleen metastatic burden were higher in the mice that were orthotopically injected with *TOB1-AS1* overexpression PANC-1 cells. The bar plots show quantified total radiance with a set area (in p/sec). **F**, Targeting *TOB1-AS1* by two ASOs. **G**, *TOB1-AS1* knockdown in Panc 10.05 and PATU-8988S cells transduced with ASO1 (n=3), ASO2 (n=3) or non-targeting control ASO (NC) (n=3). **H**, *TOB1-AS1* knockdown suppressed Panc 10.05 (3 biological replicates) and PATU-8988S (3 biological replicates) cell invasion in vitro. Cell invasion fold change calculation is the same as in **B**. Two-sided student t test was used. Error bars in all panels indicate standard error of the mean.

It is possible that *TOB1-AS1*, as an anti-sense lncRNA, transcriptionally regulates the expression of the sense protein-coding gene *TOB1*. However, we did not find consistent correlations between *TOB1-AS1* and *TOB1* expression in different pancreatic cancer cohorts and pancreatic cancer cell lines (**Supplementary Fig. S12D**). Hence, it is unlikely that *TOB1-AS1* functions through transcriptional regulation of *TOB1*. Although knocking down *TOB1-AS1* resulted in down regulation of *TOB1* expression, this is an expected result given that the ASOs also targeted the introns of *TOB1* (**Fig. 6F**). The decrease in *TOB1* expression was relatively mild at 10-20% (**Fig. 6G**). Overexpression of *TOB1-AS1* did not have a major impact on *TOB1* expression (**Fig. 6A**). Therefore, the oncogenic functions of *TOB1-AS1* that we observed in vitro and in vivo are likely independent of *TOB1*. To gain further insights into the pathway that *TOB1-AS1* is involved in and its downstream targets, we performed RNA-Seq on PANC-1-generated mouse tumors with *TOB1-AS1* overexpression and found that the most significantly differentially expressed gene was *CNNM1* (**Supplementary Fig. S12E**). *CNNM1* is a cyclin and CBS domain divalent metal cation transport mediator and is predicted to be involved in ion transport ^52^. How *TOB1-AS1* promotes cell invasion and tumor metastasis and whether *CNNM1* plays a role require further study.

Our results showed that the lncRNA *TOB1-AS1* is oncogenic and has a pro-metastatic function in pancreatic cancer, and that HYENA is able to detect novel proto-oncogenes activated by distal enhancers.

## Discussion

Here, we report a computational algorithm HYENA to detect candidate oncogenes activated by distal enhancers via somatic SVs. These SV breakpoints fell in the regulatory regions of the genome and caused a shuffling of regulatory elements, altering gene expression. The candidate genes we detected were not limited to protein-coding genes but also included non-coding genes. Our in vitro and in vivo experiments showed that a lncRNA identified by HYENA, *TOB1-AS1*, was a potent oncogene in pancreatic cancers.

HYENA detects candidate genes based on patient cohorts rather than individual samples. Genes need to be recurrently rearranged in the cohort to be detectable, and HYENA aims to identify oncogenes recurrently activated by somatic SVs since these events are under positive selection. Therefore, sample size is a major limiting factor. Of the eight ground truth cases, HYENA only detected four (**Fig. 2A**); undetected genes were likely due to the small sample size. However, genes detected in individual tumors by tools such as cis-X and NeoLoopFinder may not be oncogenes, and recurrent events would be required to identify candidate oncogenes.

The candidate genes identified by HYENA have statistically significant associations between nearby somatic SVs and elevated expression. However, the relationship may not be causal. It is possible that the presence of SVs and gene expression are unrelated, but both are associated with another factor. We modeled other factors to the best of our ability including gene dosage, tumor purity, patient sex, age, and principal components of gene expression. In addition, it is also possible that the high gene expression caused somatic SVs. Open chromatin and double helix regions unwound during transcription are prone to double-strand DNA breaks which may produce somatic SVs. Therefore, it is possible that some of the candidate genes are not oncogenes. Functional studies are required to determine the disease relevance of the candidate genes.

Note that the predicted 3D genome organization is not cell-type-specific. Akita was trained on five high quality Hi-C and Micro-C datasets (HFF, H1hESC, GM12878, IMR90 and HCT116) ^23^ and predicts limited cell-type-specific differences. Therefore, the predicted TADs reflect conserved 3D genome structure in the five cell types (foreskin fibroblast, embryonic stem cell,

B-lymphocyte, lung fibroblast and colon cancer). There were minor differences between HFF and H1hESC (**Supplementary Fig. S3**) in genome organization. For example, the left boundary of the TAD at the *UQCRFS1* locus was different between HFF and H1hESC (**Supplementary Fig. S3A**). Nonetheless, the translocation between chromosomes 17 and 19 removed the left boundary and merged the right side of the *UQCRFS1* TAD with the *TOB1-AS1* TAD (**Fig. 4D**). Therefore, the cell-type difference likely does not have a major impact on our results.

## Acknowledgements

We thank the Center for Research Informatics at the University of Chicago for providing the computing infrastructure, Matthew Stephens for helpful suggestions, Marsha Rosner for assistance in lentiviral experiments, and Ani Solanki for assistance in animal experiments. The work was supported by the Goldblatt Endowment (A.Y.), the National Institutes of Health grant K22CA193848 (L.Y.), R01CA269977 (L.Y.) and University of Chicago and UChicago Comprehensive Cancer Center (L.Y.).

## Disclosure

The authors have no competing interests to declare.

## Notes

### Competing Interest Statement

The authors have declared no competing interest.

### Summary of Updates

Modified the HYENA algorithm and updated results.

