## Supplementary figures for "HYENA detects oncogenes activated by distal enhancers in cancer"

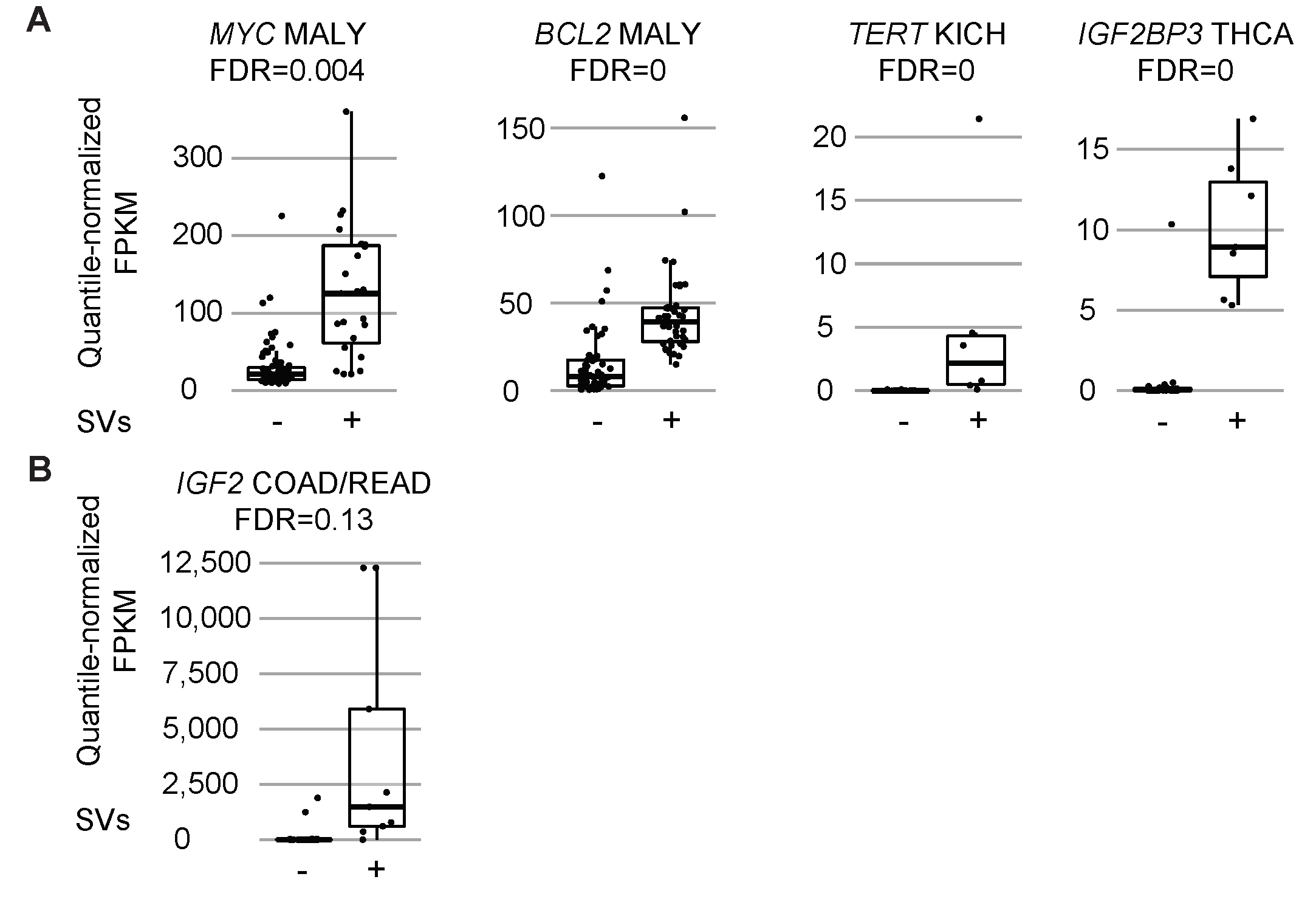

**Supplementary Fig. S1. Expression levels for five known enhancer hijacking target oncogenes.** For each gene, tumors are grouped based on SV status (- or +). Quantile normalized FPKM values are shown for each group. The boxplots show median values (thick black lines), upper and lower quartiles (boxes), and 1.5× interquartile range (whiskers). Individual tumors are shown as black dots. **A**, Genes detected by HYENA. **B**, Gene not detected by HYENA.

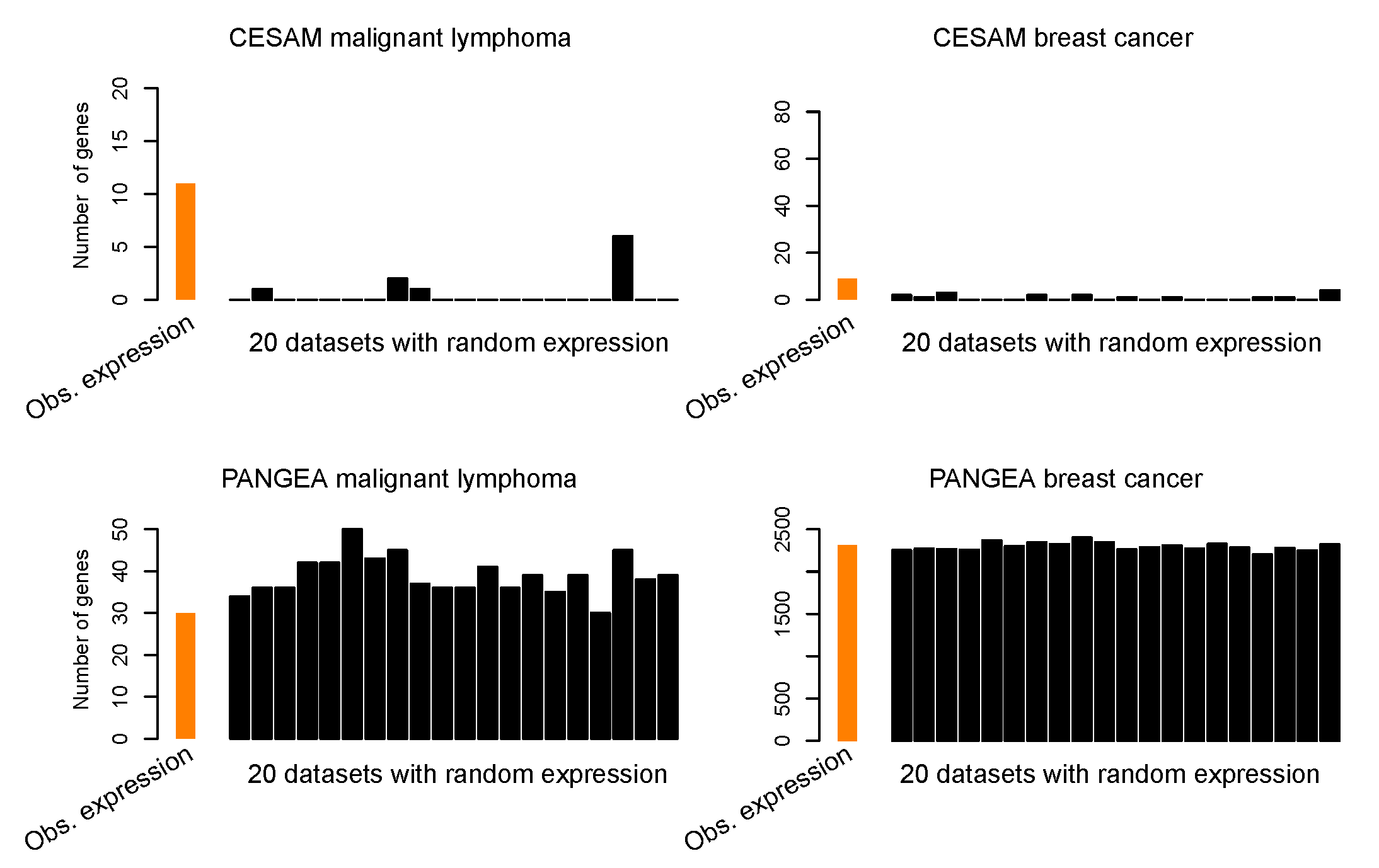

**Figure S2. Numbers of genes detected by CESAM and PANGEA in two PCAWG tumor types using observed gene expression and randomized expression.** Genes detected when expression was randomized were false positives.

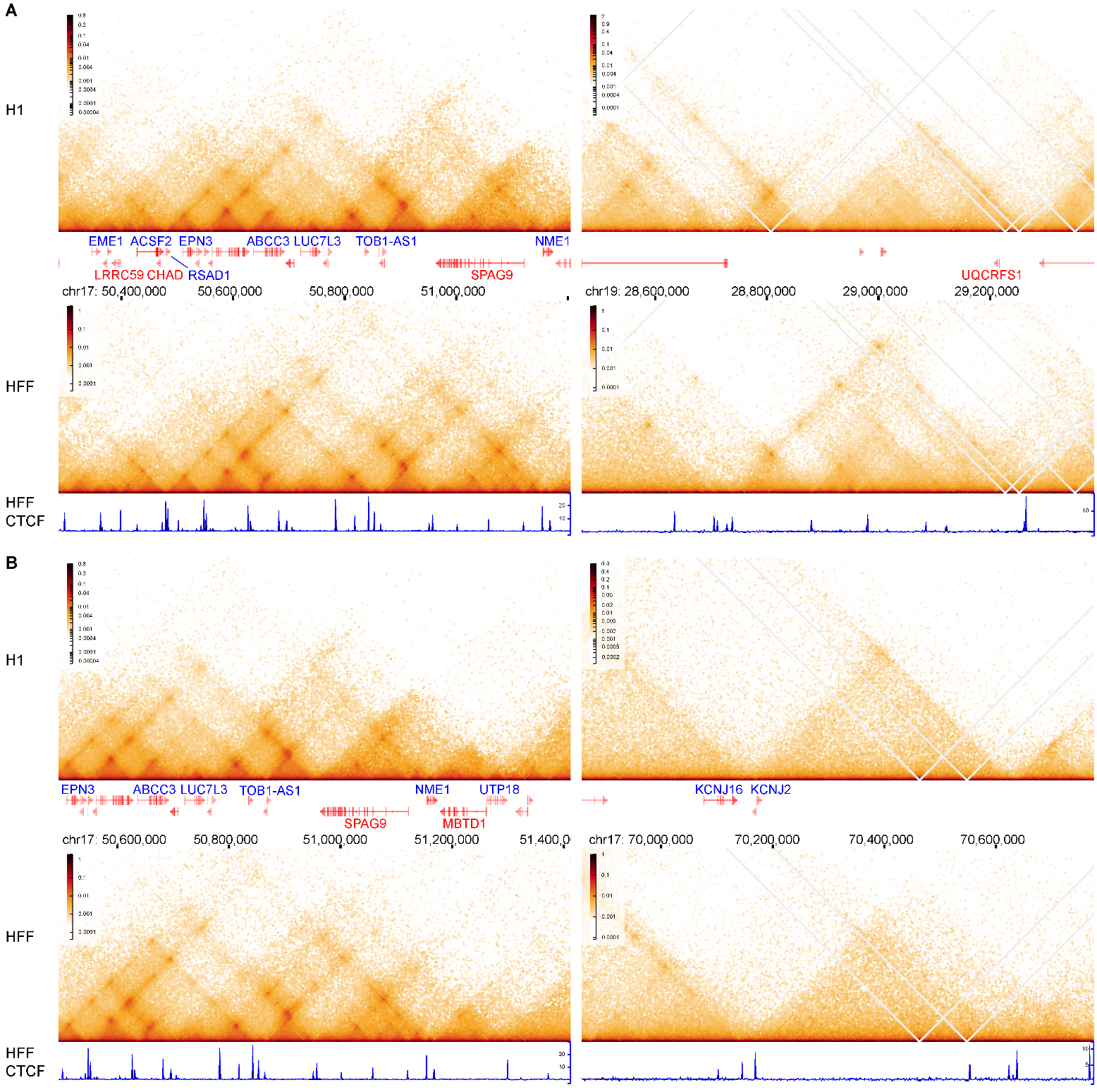

**Figure S3.** **Hi-C maps of *TOB1-AS1*, *UQCRFS1*, and *KCNJ2* loci from H1 and HFF cell lines.** **A**, *TOB1-AS1* (left panels) and *UQCRFS1* (right panels) loci. **B**, *TOB1-AS1* (left panels) and *KCNJ2* (right panels) loci. CTCF ChIP-seq of the HFF cell line is shown at the bottom. These experiment-based Hi-C maps are very similar to predicted Hi-C maps for the same loci in **Fig. 4D** and **4E** left and middle panels.

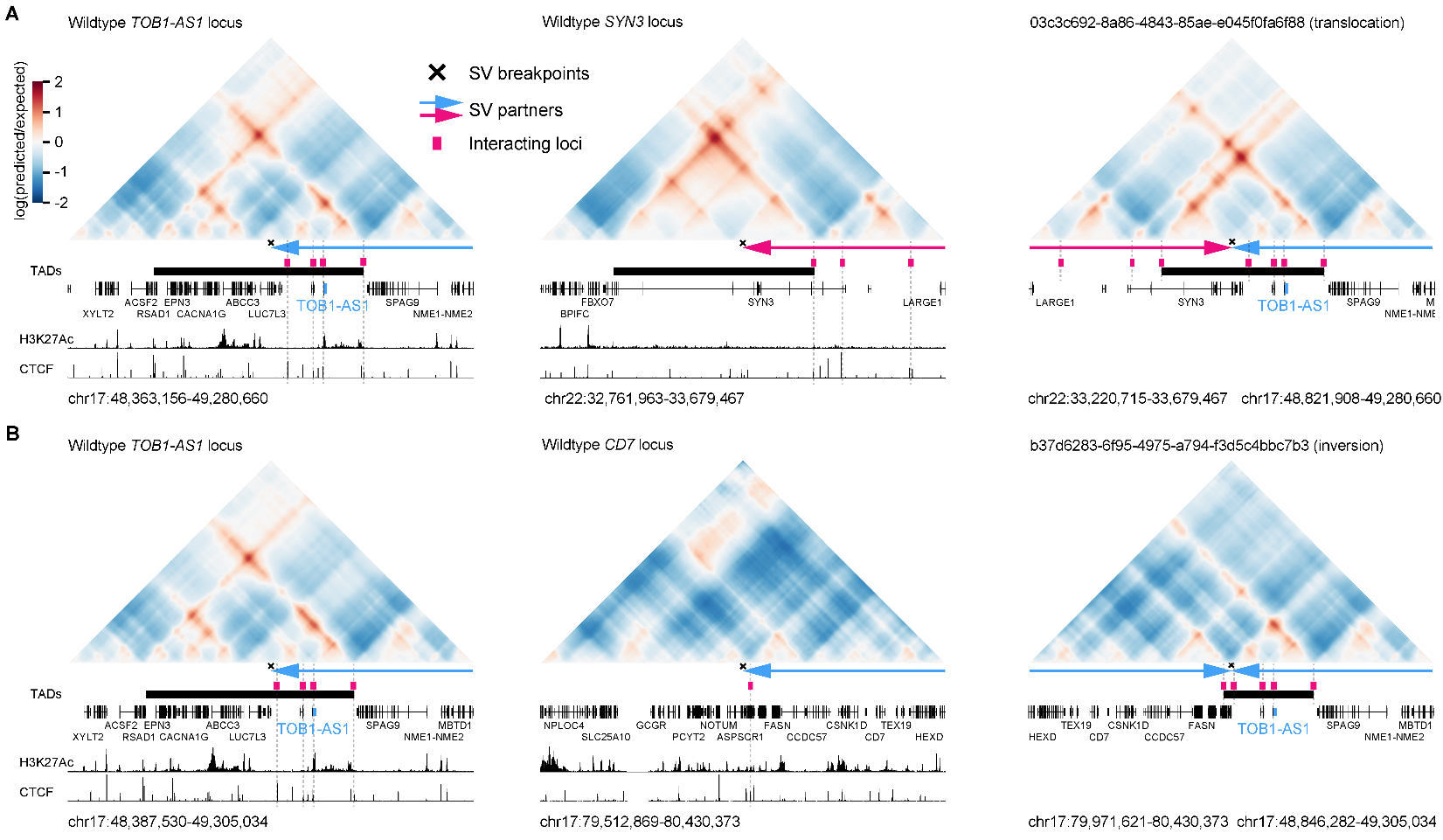

**Figure S4. Predicted 3D chromatin interaction maps for two pancreatic cancers with SVs near *TOB1-AS1*. A**, Predicted maps for regions without translocations (left and middle panels) and with translocation in tumor 03c3c692-8a86-4843-85ae-e045f0fa6f88 (right panel). **B**, Predicted maps for regions without inversion (left and middle panels) and with inversion in tumor b37d6283-6f95-4975-a794-f3d5c4bbc7b3 (right panel).

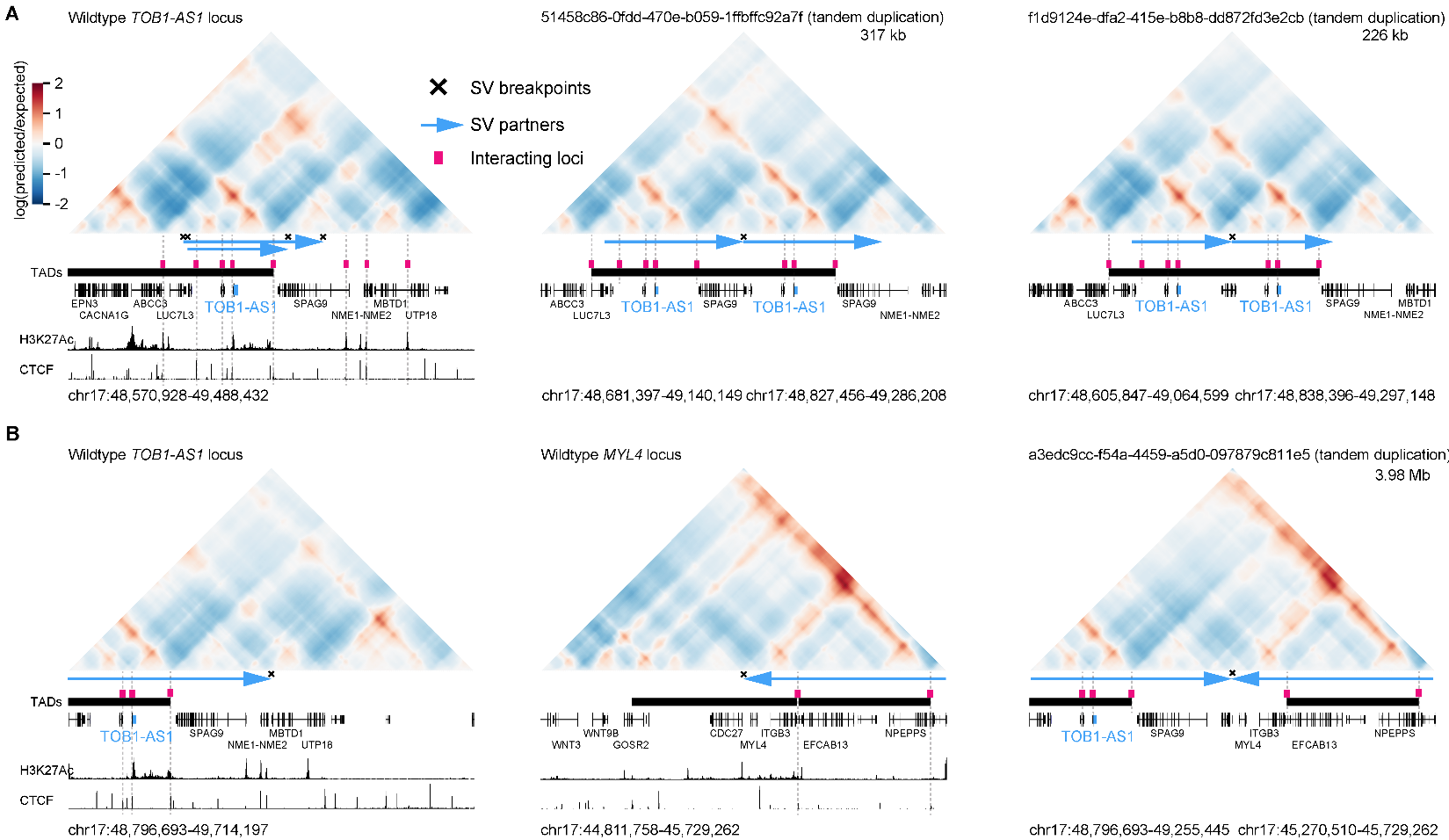

**Figure S5. Predicted 3D chromatin interaction maps for three pancreatic cancers with SVs near *TOB1-AS1*. A**, Predicted maps for regions without tandem duplication (left panel) and with tandem duplications in two tumors 51458c86-0fdd-470e-b059-1ffbffc92a7f (middle panel) and f1d9124e-dfa2-415e-b8b8-dd872fd3e2cb (right panel). **B**, Predicted maps for regions without tandem duplication (left and middle panels) and with tandem duplication in tumor a3edc9cc-f54a-4459-a5d0-097879c811e5 (right panel).

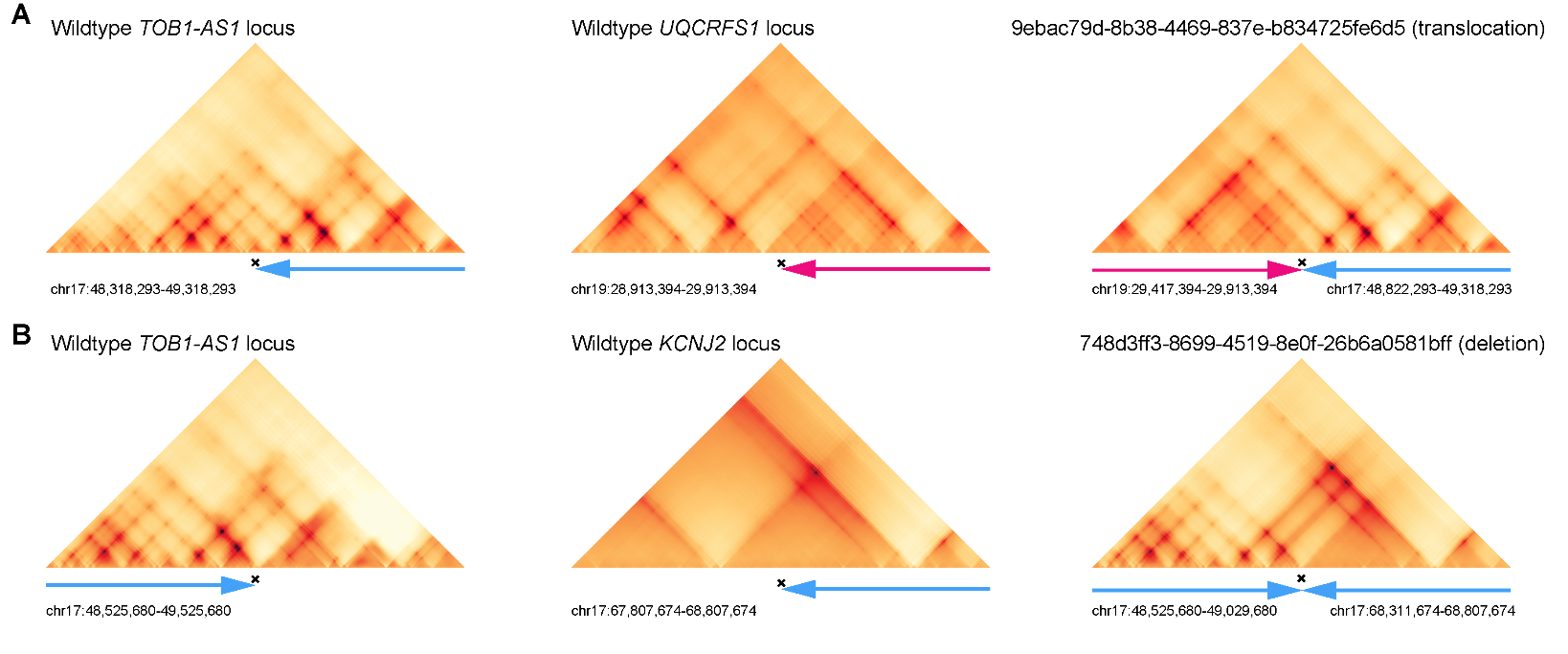

**Figure S6. 3D genome structures predicted by deep-learning based algorithm Orca.** **A**, Predicted 3D chromatin interaction maps of the *TOB1-AS1* (left panel), *UQCRFS1* (middle panel), and the translocated region in tumor 9ebac79d-8b38-4469-837e-b834725fe6d5 (right panel). **B**, Predicted 3D chromatin interaction maps of *TOB1-AS1* (left panel) and *KCNJ2* (middle panel) loci without deletion as well as the region after deletion in tumor 748d3ff3-8699-4519-8e0f-26b6a0581bff (right panel). The 6 regions in this figure are the same regions shown in **Fig. 4D** and **4E**.

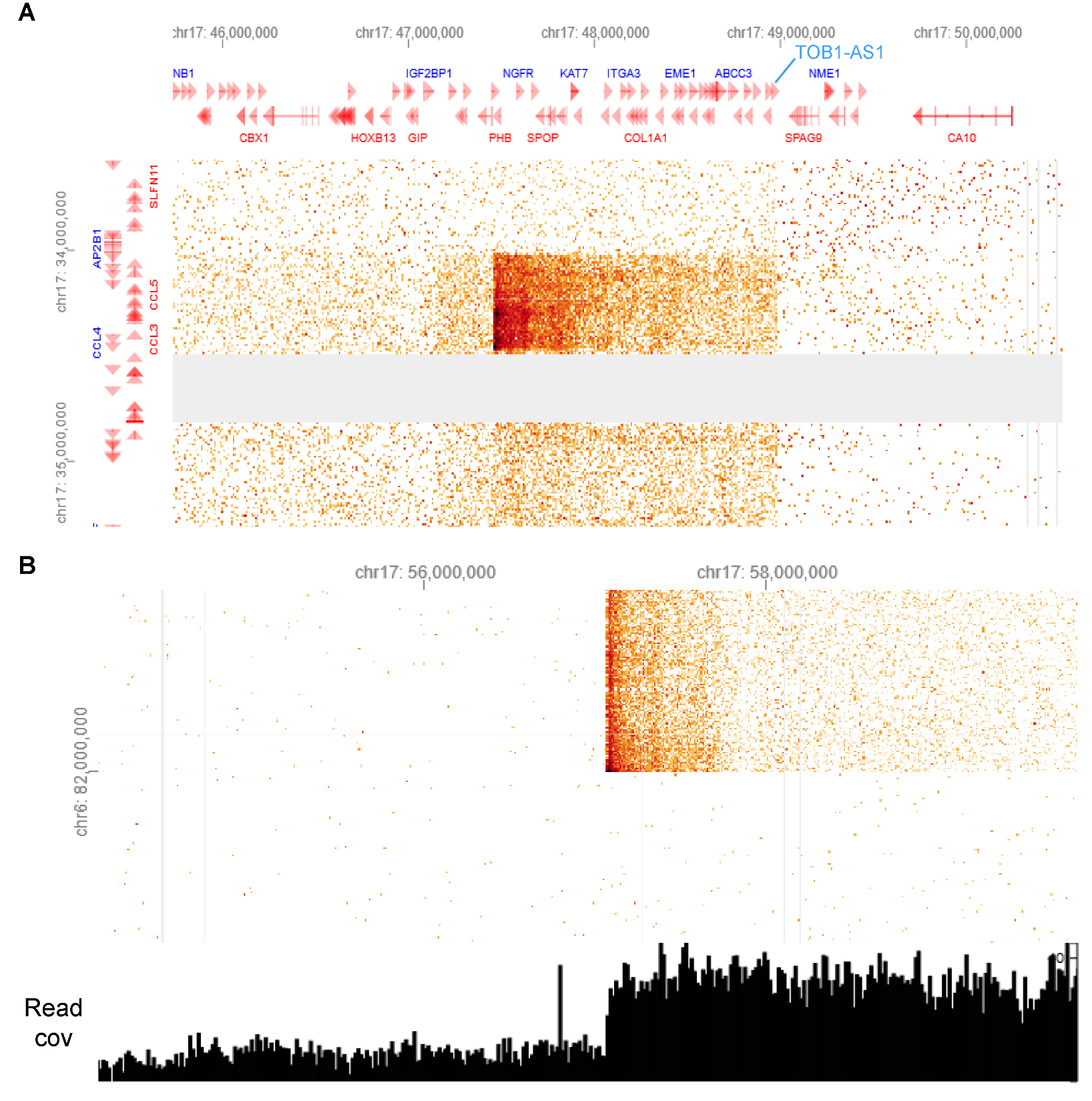

**Figure S7. SVs in Panc 10.05 detected by Hi-C.** **A**, HiGlass view showing a deletion of chr17:34,460,000-47,450,000. **B**, HiGlass view showing a translocation between chromosomes 6 and 17. Read coverage is shown below the Hi-C contact map. The chromosome 17 translocation breakpoint is 8 Mb downstream of the CNV breakpoint shown in **Fig. 5C** left most panel.

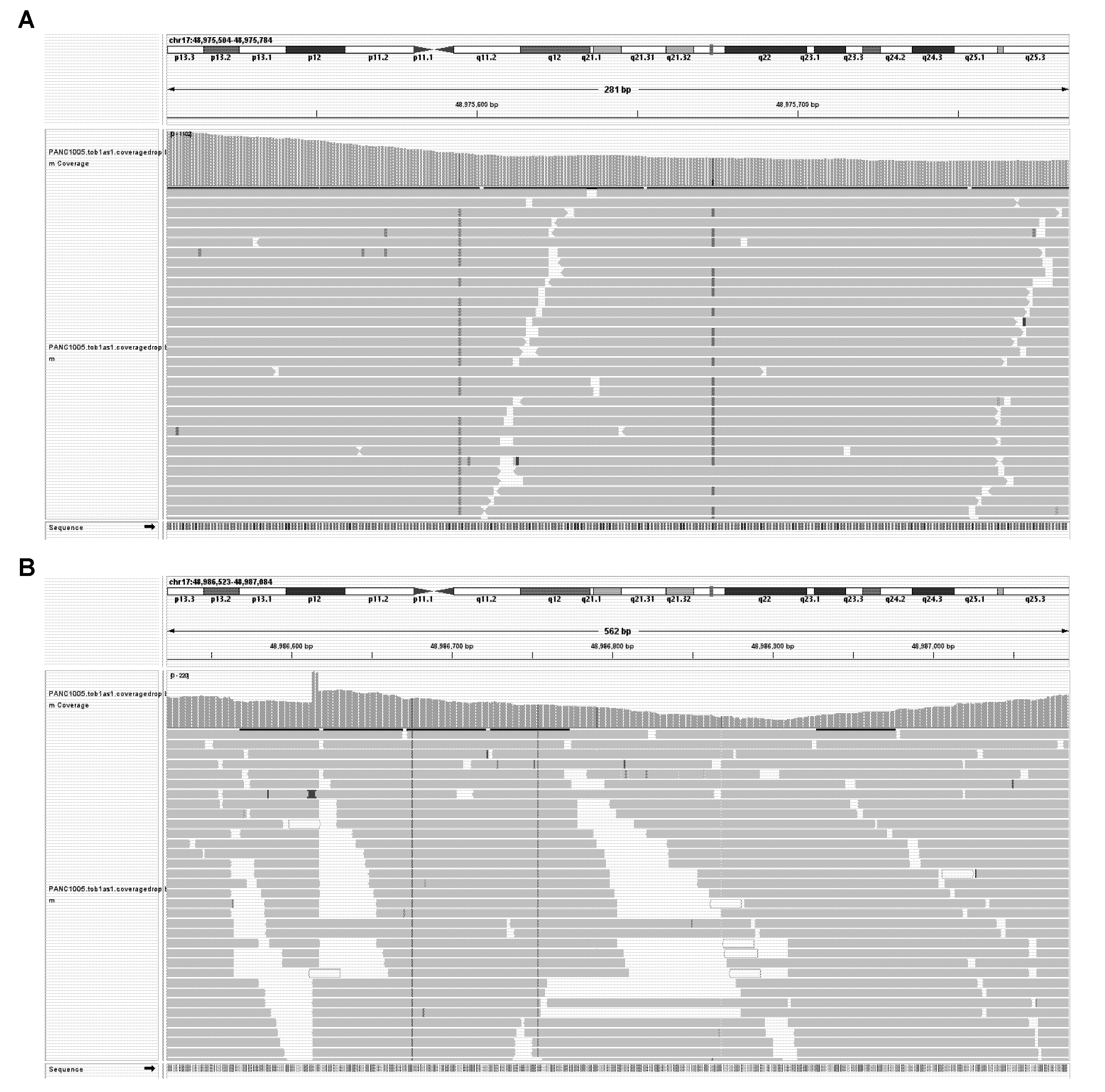

**Figure S8. SNPs in Panc 10.05 near CNV and foldback inversion breakpoint.** **A** and **B**, IGV screenshots showing reads mapped to five-copy and one-copy regions in Panc 10.05 in **Fig. 5C** left most panel. Horizontal grey bars are Hi-C sequencing reads. Colored lines are mismatches of reads compared to the reference genome. Grey vertical bars are read depth. Colored vertical bars represent SNPs. The two SNPs in **A** are heterozygous SNPs, whereas the four in **B** are homozygous.

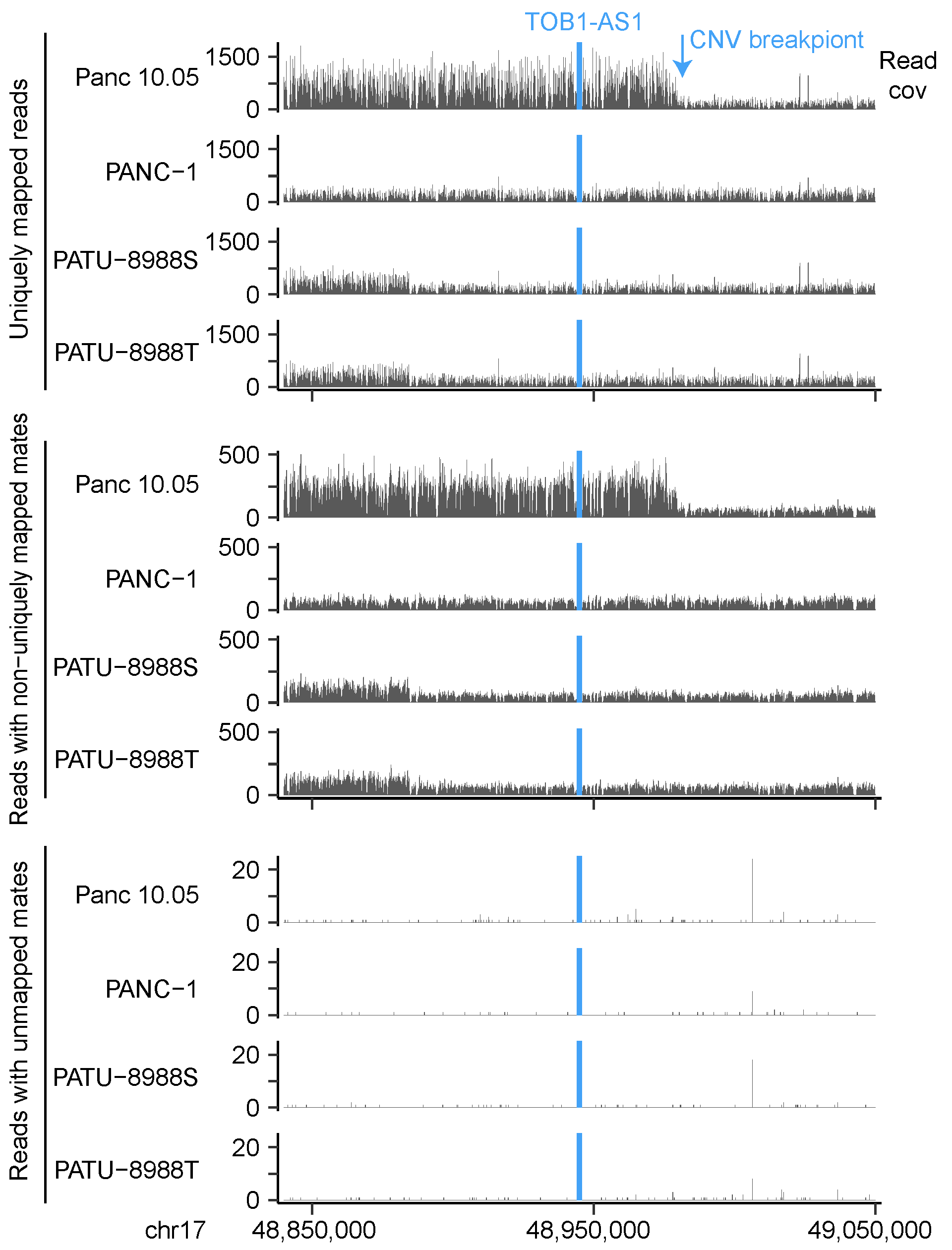

**Figure S9. Read coverage in 4 pancreatic cell lines at the *TOB1-AS1* locus.** The numbers of uniquely mapped reads, reads with non-uniquely mapped mates, and reads with unmapped mates are shown for four pancreatic cancer cell lines. In Panc 10.05, the ratio of non-uniquely mapped mates in the high-copy region (upstream of the CNV breakpoint) and the low-copy region (downstream of the CNV breakpoint) is comparable to that of uniquely mapped reads. Very few unmapped mates are present in any cell lines.

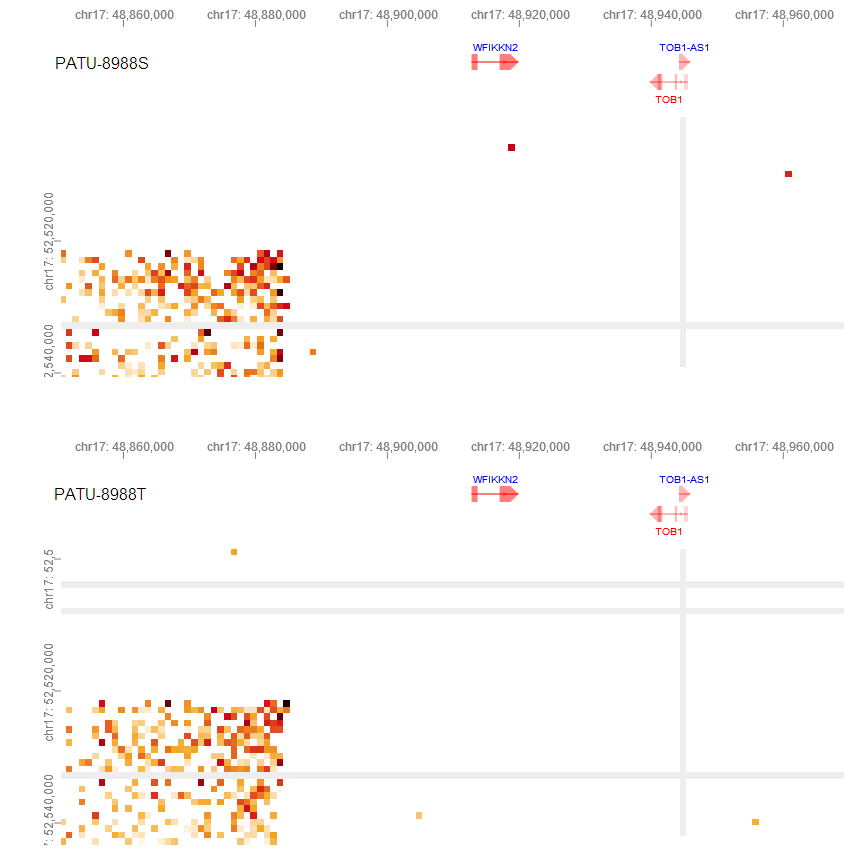

**Figure S10. HiGlass views showing the shared SV near *TOB1-AS1* in PATU-8988S (top) and PATU-8988T (bottom).** The SV is about 50 kb upstream of *TOB1-AS1* and points away from *TOB1-AS1*. The locations of *TOB1-AS1* are shown in the x-axis at the top.

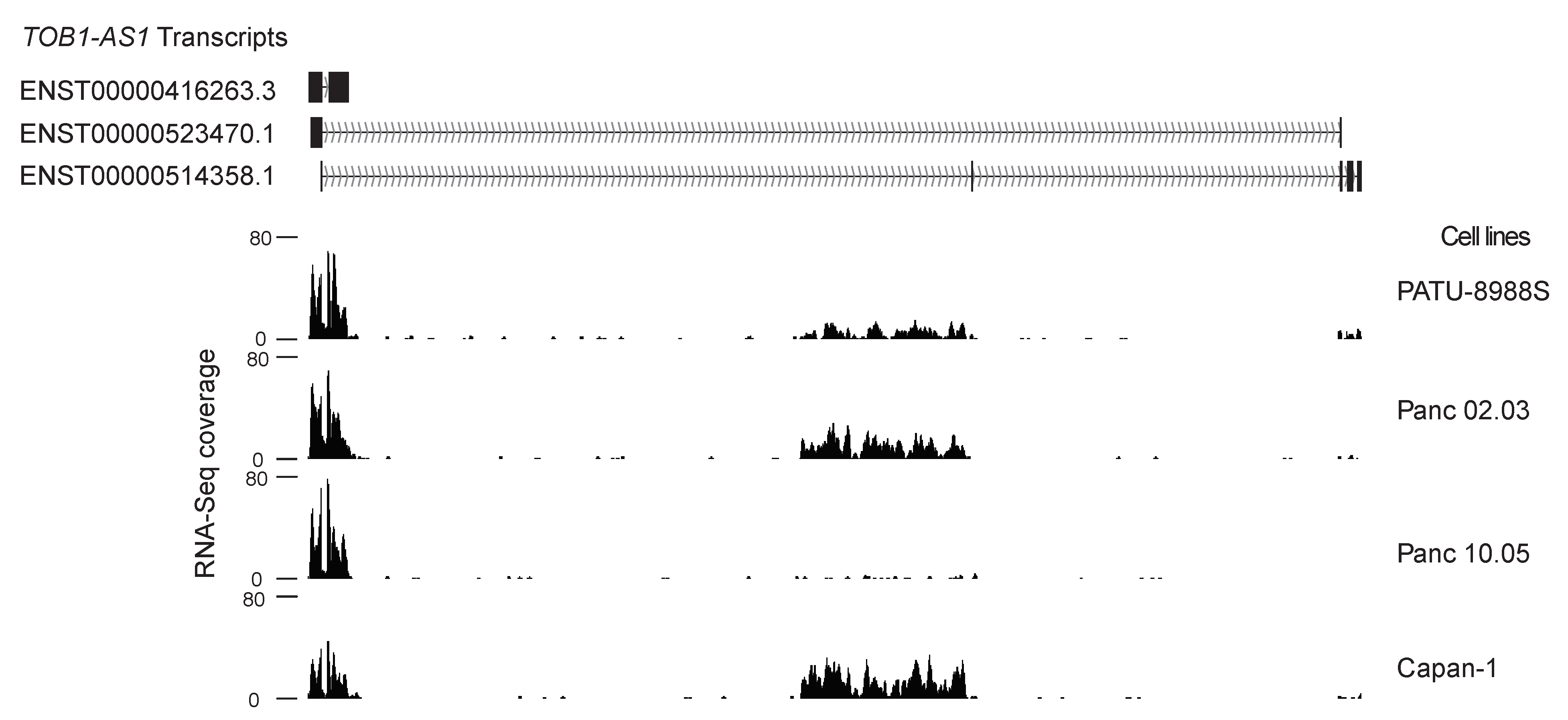

**Figure S11. RNA-Seq coverage of *TOB1-AS1* isoforms.** RNA-Seq coverage of three *TOB1-AS1* isoforms from four pancreatic cancer cell lines with high *TOB1-AS1* expression (PATU-8988S, Panc 02.03, Panc 10.05, and Capan-1). The major isoform is ENST00000416263.3.

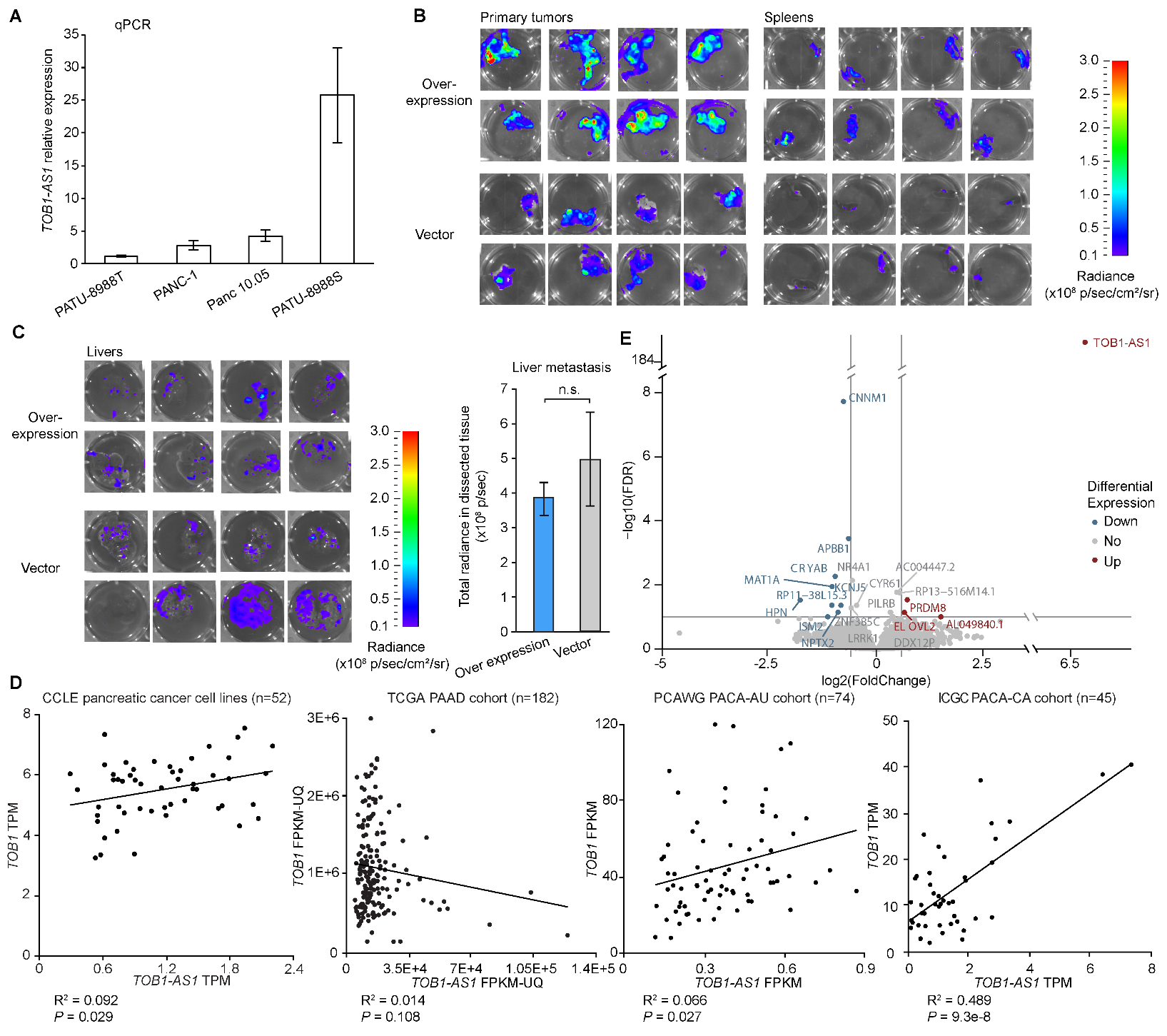

**Figure S12. *TOB1-AS1* overexpression.** **A**, *TOB1-AS1* relative expression levels in PATU-8988T, PANC-1, Panc 10.05, and PATU-8988S cell lines based on quantitative RT-PCR. The relative expression of the other three cell lines was calculated relative to PATU-8988T. Error bars indicate standard error of the mean. **B**, Ex vivo IVIS images showing primary tumors and spleen metastatic tumors from mice orthotopically injected with PANC-1. **C**, Ex vivo IVIS images and radiance quantification (p/sec) of whole wells showing liver metastatic tumors in mice orthotopically injected with PANC-1. Two-sided student t test was used. Error bars indicate the standard error of the mean. **D**, Scatter plots showing the correlations between *TOB1* and *TOB1-AS1* RNA expression in CCLE pancreatic cancer cell lines, TCGA PAAD, PCAWG PACA-AU, and ICGC PACA-CA cohorts. Sample sizes, gene expression normalization methods, squared-Rs and *P* values are labeled. In CCLE cell lines and the PCAWG PACA-AU cohort, the two genes have very weak positive associations with marginal *P* values of 0.029 and 0.027. In the ICGC PACA-CA cohort, the two genes have a strong positive correlation. However, the correlation is mainly driven by two outliers. On the contrary, in the TCGA PAAD cohort, the two genes are not significantly correlated. Therefore, *TOB1-AS1* and *TOB1* do not have consistent associations in patient samples and cell lines. **E**, Volcano plot showing the differentially expressed genes in *TOB1-AS1* overexpression PANC-1 tumors (n=6) compared to vector control tumors (n=6). Red and blue dots with gene labels represent significantly (FDR <0.1) upregulated and downregulated genes with fold-change larger than 1.5 and smaller than 1/1.5, respectively. Grey dots represent all other genes. Grey lines represent -log10(FDR) of 1 (horizontal), log2(FoldChange) of log2(1.5) (vertical, right) and log2(1/1.5) (vertical, left).
